# Growth Factor Independence (GFI) 1B-mediated transcriptional repression and lineage allocation require Lysine Specific Demethylase (LSD)1-dependent recruitment of the BHC complex

**DOI:** 10.1101/519090

**Authors:** David McClellan, Mattie J. Casey, Diana Bareyan, Helena Lucente, Christopher Ours, Matthew Velinder, Jason Singer, Mehraju Din Lone, Wenxiang Sun, Yunuen Coria, Clinton Mason, Michael E. Engel

## Abstract

Growth Factor Independence (GFI)1B coordinates assembly of transcriptional repressor complexes comprised of co-repressors and histone modifying enzymes to control gene expression programs governing lineage allocation in hematopoiesis. Enforced expression of GFI1B in K562 erythroleukemia cells favors erythroid over megakaryocytic differentiation, providing a platform to define molecular determinants of binary fate decisions triggered by GFI1B. We deployed proteome-wide proximity labeling to identify factors whose inclusion in GFI1B complexes depends upon GFI1B’s obligate effector, Lysine Specific Demethylase (LSD)1. We show that GFI1B preferentially recruits core and putative elements of the BRAF-histone deacetylase (HDAC) (BHC) chromatin remodeling complex (LSD1, RCOR1, HMG20A, HMG20B, HDAC1, HDAC2, PHF21A, GSE1, ZMYM2 and ZNF217) in an LSD1-dependent manner to control erythroid fate specification. Among these, depletion of both HMG20A and HMG20B, or GSE1 block GFI1B-mediated erythroid differentiation, phenocopying impaired differentiation brought on by LSD1 depletion or disruption of GFI1B—LSD1 binding. These findings demonstrate the central role of the GFI1B—LSD1 interaction as a determinant of BHC complex recruitment to enable cell fate decisions driven by GFI1B.

## INTRODUCTION

Transcriptional regulation is critical for cell growth, homeostasis, and fate determination in multicellular organisms. In hematopoiesis, cell surface receptors, signaling networks, transcription factors and their effectors cooperate to influence chromatin state and gene expression, directing hematopoietic progenitors toward specific fates, and once achieved, ensuring proper function for each cell type. Failure to properly control chromatin structural dynamics governing stage-specific gene expression programs can cause qualitative and quantitative hematopoietic disorders, including malignancies. Failures can arise from altered expression of transcription factors or from malfunction or misdirection of their partners (1–3). Thus, knowing the composition of protein complexes that regulate gene expression by cell fate specifying transcription factors is essential to understanding normal and malignant hematopoiesis, and may have therapeutic relevance (4).

Growth Factor Independence (GFI) family transcriptional repressors, GFI1 and GFI1B, are master regulators of developmental hematopoiesis (5, 6). GFI1 controls hematopoietic stem cell self-renewal, B-and T-lymphopoiesis, and directs the granulocyte versus monocyte fate decision during late stages of myeloid differentiation (7–9). Notably, *Gfi1-null* mice are viable, but are characterized by severe neutropenia with compensatory monocytosis and sensorineural hearing loss (10–12). Altered *GFI1* expression is found in multiple hematologic malignancies, most notably acute myeloid leukemia (AML) and T-cell acute lymphoblastic leukemia (T-ALL) (13–17). Moreover, loss of function mutations in *GFI1* cause severe congenital neutropenia (SCN) type 2 and nonimmune chronic idiopathic neutropenia (CIN) of adults, both of which predispose to AML (18, 19). GFI1B is required for definitive erythropoiesis and megakaryopoiesis, and like GFI1, also helps to maintain hematopoietic stem cell quiescence (20–23). In mice, *Gfi1b* nullizygosity is lethal at embryonic day (E) 15 due to erythropoietic failure (24). Mutations in *GFI1B* are associated with both qualitative and quantitative disorders of platelets and elevated *GFI1B* expression is found in AML (25–28). However, the role of GFI1B in AML is uncertain. Despite *GFI1B* often being increased in AML patients (29–32), a recent study suggests low *GFI1B* expression correlates with poor patient outcome and that in mice, *Gfi1b* depletion accelerates progression to AML from myelodysplasia driven by a NUP98-HOXD13 fusion protein (33). Collectively, these observations suggest widespread contributions by GFI family proteins to lineage allocation during normal hematopoiesis and both oncogenic and oncosuppressor roles in hematologic malignancies. A clear understanding of GFI1/1B contributions to hematopoiesis and malignancy requires comprehensive accounting of their protein partners and their dependency relationships during cell fate decision making.

GFI family proteins feature a shared structural organization (5, 34). Both GFI1 and GFI1B are comprised of a Snail/Slug/Gfi1 (SNAG) domain and a highly conserved concatemer of six C2H2-type zinc-fingers (ZnF) at their N-and C-termini, respectively. Linker regions with limited conservation separate the SNAG and ZnF domains. Sequence-specific DNA binding by both GFI1 and GFI1B is coordinated by ZnFs 3, 4 and 5 (32, 35–37), while ZnFs 1, 2, and 6 enable protein—protein interactions with an incompletely defined subset of transcriptional partners (38–41). Linker regions provide a platform for protein binding and post-translational modifications that regulate innate functions of GFI proteins (40, 42). GFI1 and GFI1B SNAG domains differ only by a conservative S→T substitution at amino acid 14, but are otherwise identical. Moreover, the SNAG domain is near invariant among orthologs from human to zebrafish, underscoring the critical role of the SNAG domain and its binding partners. Prominent among these is Lysine Specific Demethylase (LSD)1 (KDM1A, AOF2, BHC110, NPAO, KIAA0601), which is required for transcriptional repression by GFI proteins (10, 43). The SNAG domain is necessary and sufficient for LSD1 binding by GFI1 and GFI1B (44). SNAG domain deletion, alanine substitution at proline 2 (P2A) or leucine substitution at lysine 8 (K8L) of the SNAG domain abolishes LSD1 binding to GFI family proteins and profoundly impairs their activity in multiple functional assays (43, 45–47).

LSD1 is a flavin-dependent monoamine oxidase that demethylates mono-and di-methylated histone 3 lysines 4 and 9 (H3K4me1/2 and H3K9me1/2) to impact chromatin structure and gene expression (48, 49). LSD1 also demethylates non-histone targets including p53, STAT3, ERα, MYPT1, HIF1α, DNMT1, E2F and SOX2, suggesting broad influence over cellular homeostasis (50–57). LSD1 operates within multiple assemblies that regulate transcription, including <u>Nu</u>cleosome <u>R</u>emodeling and <u>D</u>eacetylase (NuRD), <u>C-t</u>erminal <u>B</u>inding <u>P</u>rotein (CtBP), <u>M</u>ixed <u>L</u>ineage Leukemia (MLL) coactivator and BRAF-Histone deacetylase (BHC) complexes (58–62). Its inclusion in distinct multiprotein assemblies suggests both enzymatic and structural contributions to transcriptional control, yet affiliations between LSD1-containing complexes and the DNA binding proteins that recruit them are poorly understood.

We leveraged GFI1B-driven erythroid differentiation in K562 cells, GFI1B fusion proteins with a promiscuous biotin ligase (BirA*) (63, 64) and unbiased mass-spectral quantitation to define proximity relationships between GFI1B and partners that depend upon GFI1B—LSD1 binding. We show that GFI1B—LSD1 binding is necessary but not sufficient for GFI1B-mediated erythroid differentiation in K562 cells. We also show that recruitment of core (65) (LSD1, RCOR1, HDAC1, HDAC2, HMG20B and PHF21A) and putative (66, 67) (HMG20A, GSE1, ZMYM2, and ZNF217) BHC complex components by GFI1B depends upon GFI1B—LSD1 binding and that depletion of candidate BHC complex proteins HMG20A and HMG20B, as well as GSE1 impairs GFI1B-mediated erythroid differentiation even though GFI1B—LSD1 interaction potential remains. Our results show that GFI1B-mediated lineage allocation depends upon LSD1’s function as a scaffolding component for BHC complex assembly and highlight the need to consider protein assemblies as modules which act in concert as determinants of cell fate.

## RESULTS

### LSD1 depletion impairs GFI1B-mediated erythroid differentiation

The human erythroleukemia cell line, K562, displays erythroid differentiation with enforced GFI1B expression (68), reflected by hemoglobinization and benzidine positive staining (Fig. 1A). Likewise, K562 cells adopt megakaryocytic features, including CD61 expression, with tetradecanoyl-1-phorbol-13-acetate (TPA) treatment (Fig. 1B). These dual fates are mutually exclusive, enabling GFI1B dependency relationships relevant to fate specification to be discerned. Notably, deletion of the GFI1B SNAG domain or placement of an epitope tag at the GFI1B N-terminus abolishes GFI1B-driven erythroid differentiation, suggesting primacy of the SNAG domain for this phenotype (68–70). Being necessary and sufficient for LSD1 binding, we surmised that disrupting or blocking access to the SNAG domain would impair LSD1 recruitment, and by extension, that LSD1 function would be required for GFI1B-driven erythroid differentiation. To confirm this, we depleted K562 cells of LSD1 and tested GFI1 B’s ability to induce erythroid differentiation and for TPA to trigger cell surface expression of CD61. Immunoblots show depletion of LSD1 by two distinct shRNA constructs relative to a scramble control sequence, and equivalent expression of GFI1B (Fig. 1C). LSD1 depletion almost completely eliminates the detection of benzidine positive cells in response to enforced expression of GFI1B compared to controls yet had limited impact on CD61 expression at the cell surface following TPA treatment (Fig. 1D and E). LSD1 depletion also blocked expression of alpha (*HBA1/2*) and beta (*HBB*) globins and alpha hemoglobin stabilizing protein (*AHSP)*, as well as 5-aminolevulinic acid synthase (*ALAS)2* and ferrochelatase (*FECH*), which are rate limiting enzymes required for heme biosynthesis. Expression of the erythroid fate marker glycophorin A (*GYPA*) was also significantly impaired (Fig. 1F and G). To confirm contributions to GFI1B-mediated transcriptional repression, we examined mRNA levels for GFI1B direct targets *CDKN1A, GFI1B* and *MYB* in the context of LSD1 depletion. Repression of each gene was reversed by LSD1 depletion (Fig. 1H). Together these data show that LSD1 enables transcriptional repression by GFI1B to trigger an erythroid fate specifying program in K562 cells and that LSD1 is dispensable for TPA-driven megakaryocytic differentiation.

**Figure 1.**
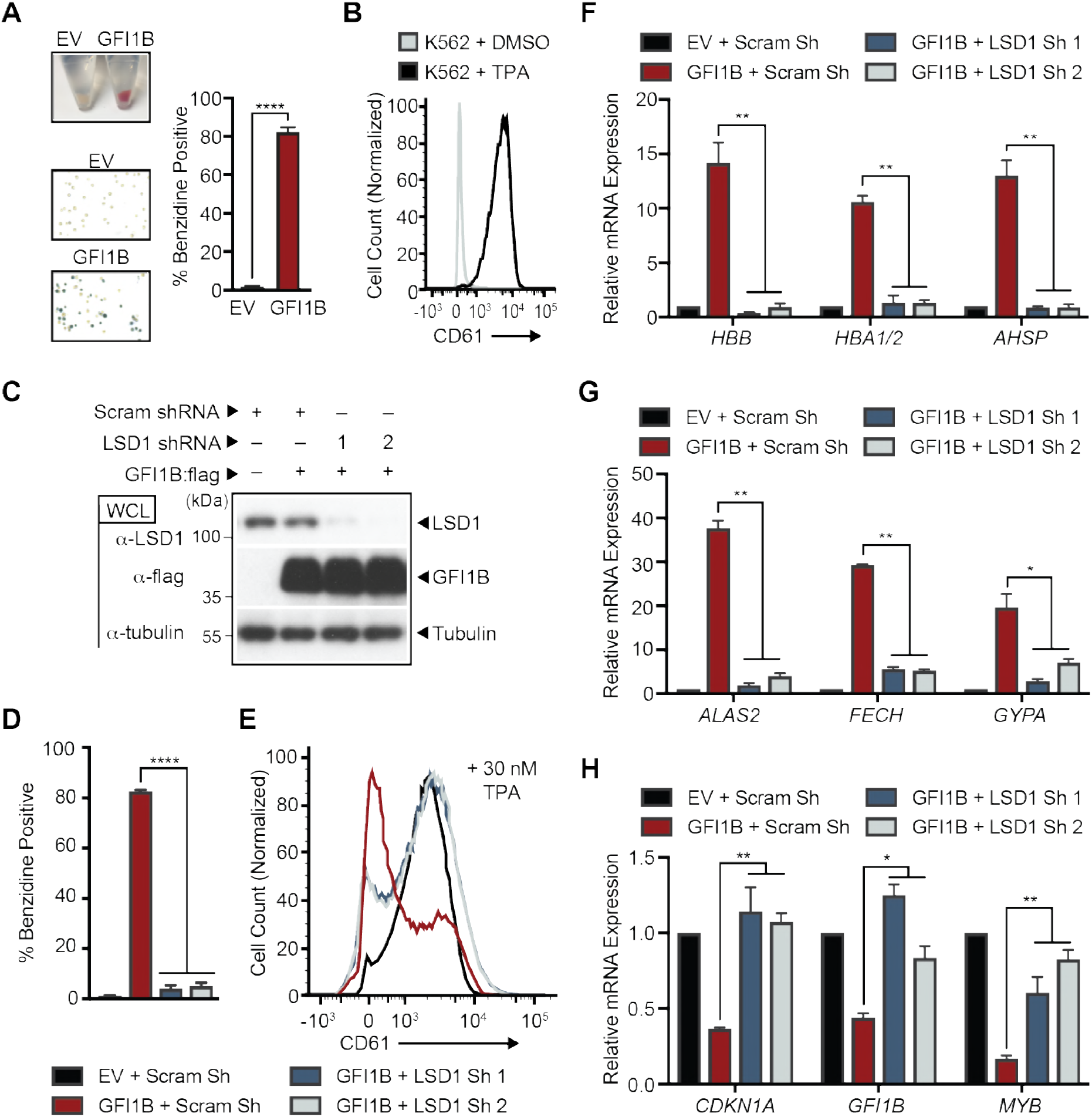
LSD1 depletion impedes GFI1B-mediated erythroid differentiation. (A) GFI1B overexpression promotes K562 cell hemoglobinization. Cells were infected with empty vector (EV) or virus expressing GFI1B. GFI1B-expressing cells become red and are benzidine positive. Benzidine positive cells are quantified as mean ± SD from four biological replicates. (B) TPA triggers cell surface expression of CD61 in K562 cells. CD61 expression was quantified in vehicle-vs. TPA-treated viable cells by flow cytometry. (C) LSD1 depletion and GFI1B overexpression in K562 cells. Small hairpin RNAs targeting *LSD1* (LSD1 shRNA) or content-matched scramble control (Scram shRNA) were inducibly transcribed in K562 cells followed by infection at day +3 with GFI1 B:flag-expressing virus or EV. LSD1 and GFI1B were quantified by western blot of whole cell lysates (WCL). Tubulin served as a loading control. (D) LSD1 depletion blocks GFI1B-mediated K562 cell hemoglobinization. (E) LSD1 depletion enables TPA-induced CD61 expression in the context of GFI1B overexpression. (F-G) LSD1 depletion impairs expression of erythroid fate genes in response to GFI1B overexpression. Alpha-and Beta-globins (*HBA1/2* and HBB), alpha globin stabilizing protein (*AHSP)*, aminolevulinic acid synthase (*ALAS)2*, ferrochelatase (*FECH*) and glycophorin (*GYP)A* expression are shown. (H) LSD1 depletion restores expression of GFI1 B-repressed target genes. *CDKN1A, GFI1B* and *MYB* expression are shown. Gene expression is quantified by RT-qPCR normalized to *GUS.* Results are expressed as mean ± 2SD from two independent experiments performed in triplicate. \**p* < 0.05; \*\**p* < 0.005; ***p <0.0005, \*\*\*\**p* < 0.00005.

### GFI1B—LSD1 binding is required for GFI1B-mediated erythroid differentiation

LSD1 cooperates with an extensive roster of DNA binding proteins to influence gene expression (71), and therefore could influence this binary fate decision through partnerships with factors other than GFI1 B. To test whether an interaction between GFI1B and LSD1 is needed for GFI1B-mediated erythroid differentiation, we compared K562 cells with enforced expression of GFI1 B-WT or its -P2A derivative. The P2A substitution abolishes GFI1B—LSD1 binding, as well as transcriptional repression by fusion proteins between Gal4 and either GFI1B or the SNAG domain in isolation (Fig. 2A and B). While GFI1B-WT triggers an erythroid differentiation program in K562 cells, as reflected by hemoglobinization and erythroid-specific gene expression, GFI1 B-P2A fails to do so (Fig. 2C-F). GFI1B-WT expression also blocks TPA-induced megakaryocytic differentiation in K562 cells, yet GFI1B-P2A displays no such antagonism (Fig. 2G). Additionally, failure by GFI1B-P2A either to trigger erythroid differentiation or to block CD61 expression in response to TPA correlates with impaired transcriptional repression of direct GFI1B targets (Fig. 2H). These findings indicate LSD1 recruitment by GFI1B is necessary to direct the erythroid fate-specifying program and is not required for TPA-induced megakaryocytic differentiation.

**Figure 2.**
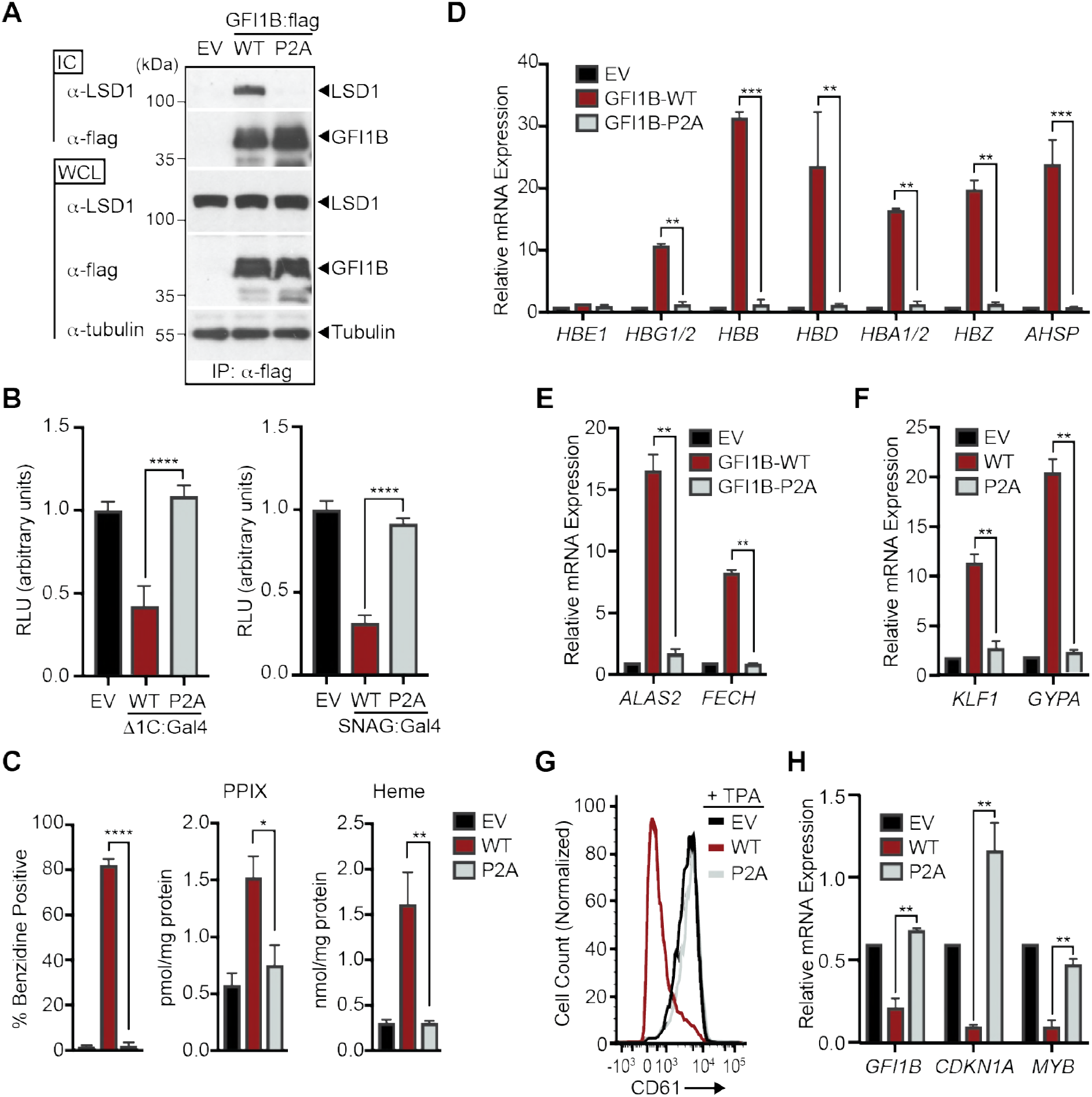
LSD1 binding is required for GFI1B-regulated cell fate decisions in K562 cells. (A) P2A substitution in GFI1B impairs GFI1B—LSD1 binding. K562 cells were transduced with constructs expressing wild type GFI1B, its P2A variant or empty vector (EV) as shown. GFI1B forms were immune purified with α-flag (M2) antibody and Protein-G Sepharose. LSD1 was quantified in immune complexes (IC) by western blot. Equal expression (LSD1 and GFI1B) and precipitation (GFI1 B) were evaluated by western blot of whole cell lysates (WCL) and ICs, respectively. Tubulin was used to confirm equal gel loading. (B) P2A substitution abolishes transcriptional repression by GFI1B. 293-T-Rex-5xGal-luciferase cells were transfected with WT or P2A variants of GFI1 B-Δ1 C:Gal4 (left) or SNAG:Gal4 (right) fusion proteins. Firefly luciferase was measured and normalized to constitutively expressed, co-transfected *Renilla* luciferase. Reporter activity is expressed as mean ±SD from two experiments performed in triplicate. (C) P2A substitution impairs GFI1B-mediated hemoglobinization. Benzidine staining was quantified as in 1A. Protoporphyrin IX (PPIX) and heme were quantified by UPLC in K562 cells transduced with EV, GFI1 B-WT or GFI1 B-P2A. (D-F) P2A substitution impairs expression of erythroid fate genes in K562 cells. Expression of β-globin cluster genes, *AHSP, ALAS2, FECH, GYPA* and Kruppel-like factor (*KLF)1* are shown. (G) P2A substitution abolishes GFI1B-mediated suppression of TPA-induced CD61 expression. Cell surface expression of CD61 in TPA treated cells expressing wild type GFI1B (WT) or its P2A variant was determined by flow cytometry. (H) P2A substitution reverses repression of GFI1B target genes *GFI1B, CDKN1A* and *MYB.* Results expressed as mean ± 2SD. *, *p* < 0.05; **, *p* < 0.005; ***, *p* < 0.0005; ****, *p* < 0.00005.

### GFI1B partners identified by proximity-dependent biotinylation

GFI1B—LSD1 binding is required for GFI1B-mediated transcriptional control and fate specifying functions in hematopoiesis (43, 45). However, LSD1 is a constituent of multiple transcriptional regulatory complexes. Which among these governs GFI1B function to direct erythroid differentiation is unknown. To more comprehensively catalogue GFI1B partners, we engineered a chimeric protein, GFI1 B-BirA*:HA, comprised of the complete GFI1B primary structure fused in frame with a promiscuous variant of the biotin ligase BirA (BirA*) from *Aquifex aeolicus* (63, 64, 72) and a C-terminal HA epitope tag under the control of a doxycycline-inducible promoter. Proteins with a proximity relationship to BirA*:HA or GFI1B-BirA*:HA are biotinylated spontaneously and can be further characterized (Fig. 3A). GFI1B-BirA*:HA or BirA*:HA were inducibly expressed in K562 cells and proteome-wide biotinylation was compared to K562 cells transduced with vector only. GFI1 B-BirA*:HA and BirA*:HA were equivalently expressed and produced a broad array of biotinylated protein substrates as reflected by streptavidin (SAv):HRP-mediated detection in whole cell extracts (Fig. 3B). Both qualitative and quantitative differences in the pattern of biotinylated proteins are apparent, suggesting preferential labeling of targets by BirA*:HA when anchored to GFI1B. In the vector control, a single 72kD band was seen, likely representing propionyl-CoA carboxylase expression in K562 cells (73). To further validate the proximity labeling technique, we confirmed enrichment for established GFI1B associated proteins, LSD1 and RCOR1, among biotinylated proteins from cells expressing GFI1B-BirA*:HA relative to those expressing non-targeted BirA*:HA. LSD1 binds directly to the GFI1/1B SNAG domain while RCOR1 engages GFI1B through LSD1 (43). LSD1 and RCOR1 were readily identified among biotinylated proteins from GFI1B-BirA*:HA-expressing cells collected on SAv-Sepharose beads, but only minimally so from cells expressing BirA*:HA or empty vector (Fig. 3C). Serving as an internal control for equal target biotinylation potential, GFI1 B-BirA*:HA and BirA*:HA were comparably modified with biotin, purified on SAv-Sepharose and detected by α-HA western blot (Fig. 3C). These findings indicate preferential biotinylation of factors known to have a proximity relationship with GFI1B. Moreover, they confirm detection of 1° (intrinsic), 2° (direct binding) and 3° (indirect binding) labeling events and validate this platform for identifying GFI1B partners proteome-wide.

**Figure 3.**
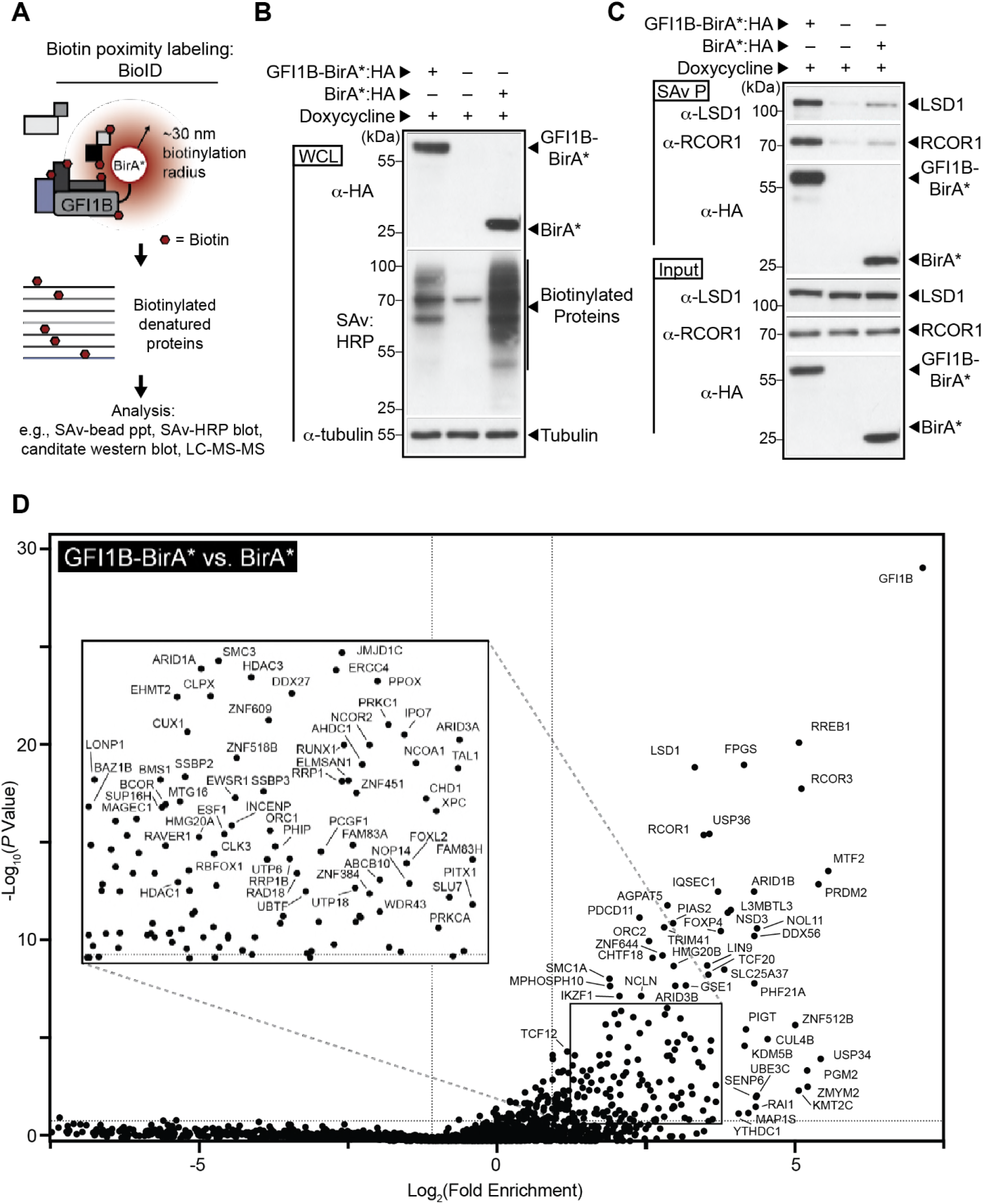
GFIB-BirA* defines a GFI1B proximitome. (A) Schematic representation of strategy for biotin modification of GFI1B partners. GFI1B is fused to the promiscuous biotin ligase with an HA epitope tag, BirA*:HA to form GFI1B-BirA*:HA. GFI1B partners are spontaneously biotinylated due to their proximity to BirA*:HA anchored to GFI1B in *cis.* Biotinylated products are captured on streptavidin (SAv)-conjugated beads for downstream analysis via candidate-based or screening approaches. (B) BirA*:HA and GFI1B-BirA*:HA fusion biotinylate diverse targets in K562 cells. Cells were transduced with constructs inducibly expressing GFI1B-BirA*:HA, BirA*:HA or EV, treated with doxycycline and harvested. GFI1B-BirA*:HA and BirA*:HA expression was confirmed by α-HA western blot and biotinylated proteins detected by SAv:HRP. Tubulin served as a loading control. (C) GFI1B interacting protein, LSD1 and RCOR1, are enriched among proteins biotinylated by GFI1B-BirA*:HA compared to BirA* or EV., Biotin-modified proteins were collected on SAv-Sepharose. LSD1, RCOR1 and the BirA* fusion proteins themselves were quantified among purified proteins (SAv P) by western blot. Equal expression was confirmed by western blot in whole cell lysates (input). (D) GFI1B-BirA*:HA identifies members of a GFI1B proximitome. Factors enriched among biotinylated and SAv-Sepharose purified proteins modified by GFI1-BirA*:HA (test) compared to BirA*:HA (control). Fold enrichment is plotted logarithmically as a ratio of average sum read intensities between test and control samples against their p-values calculated from their 20 closest ranked neighbors in the data set as described in Materials and Methods. Individual proteins are labeled with their respective gene names. Inset shows an expanded view of the data contained in the boxed area.

To comprehensively catalog GFI1B proximity partners, biotinylated proteins were collected and purified on SAv-Sepharose beads from whole cell extracts of K562 cells inducibly expressing BirA*:HA or GFI1B-BirA*:HA. Assays were performed in triplicate. Purified proteins were denatured, fractionated over acrylamide gels, excised, subjected to trypsin digestion *in situ*, then analyzed by LC-MS-MS to unequivocally establish their identities and extrapolate their abundance from sum read intensities (Supplementary Data S1). Fold enrichment (GFI1B-BirA*:HA vs. BirA*:HA only) was determined using average sum read intensities from triplicate analysis for proteins purified from GFI1B-BirA*:HA-expressing cells compared to those expressing BirA*:HA only. P-values were determined with a two-tailed Student’s t-test using variances of the 20 closest ranked proteins to account for differences in protein abundance across the data set. Fold enrichment (x-axis) for each protein is shown relative to *p*-value (*y*-axis) in a volcano plot with logarithmic axes, and statistically significant outliers are labeled (on the plot, and inset) (Fig. 3D). Notably, we observed hemoglobinization of cells expressing GFI1B-BirA*:HA comparable to those expressing GFI1B only, but not for cells expressing BirA*:HA (data not shown).

GFI1B is best known as a transcriptional repressor, and its contributions to multiple functional assays can be traced to this role (20, 74–79). Therefore, we focused first on partnerships between GFI1B and determinants of chromatin structure and transcriptional control. In the GFI1B-BirA*:HA proximitome, we identified elements of multiple complexes highly relevant to regulated gene expression (Fig. 4). Components of protein complexes enriched in the GFI1B-BirA*:HA proximitome relative to BirA*:HA control are shaded black. Those present, but not significantly above control are shaded gray and those not found in the GFI1B-BirA*:HA proximitome are shown on white background. Prominent among those proteins present and enriched in the GFI1B-BirA*:HA proximitome are members of the BHC, CtBP and BAF complexes, from which we discovered 10/10 (core + putative), 7/8 and 13/15 complex members, respectively. Similarly, we observed each component of the NuRD complex in the GFI1B-BirA*:HA proximitome, yet only a subset of these was found enriched relative to the BirA*:HA control. Components from several other complexes (NuRF, CHRAC, canonical and non-canonical PRC1, PRC2, SIN3A and NCoR) were found, and for each, some elements enriched, suggesting they also contribute to GFI1B-mediated transcriptional control and lineage allocation. Noteworthy is identification of GFI1B, being physically tethered to BirA*, as the most highly enriched protein found in this unbiased screen, further validating the strategy from a quantitative perspective.

**Figure 4.**
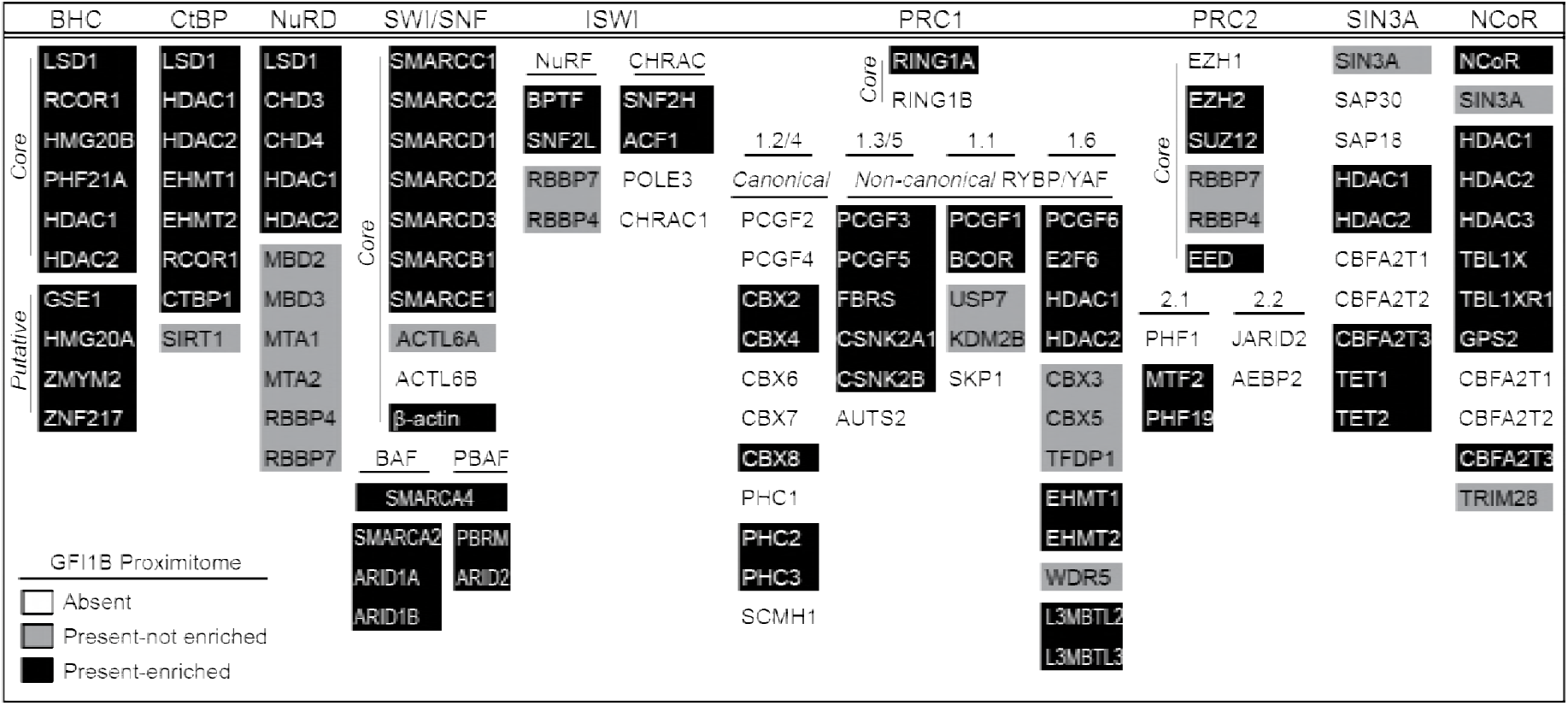
Components of transcriptional repression and chromatin remodeling complexes identified by GFI1B proximity labeling. A roster of transcriptionally relevant complexes and their component members are shown. Proteins are listed according to HUGO gene nomenclature. With respect to the GFI1 B proximitome (GFI1B-BirA*:HA vs. BirA*:HA), proteins present and enriched—white letters on black background. Those present by not significantly enriched (p <0.05)—black letters on gray background. Not present (average sum read intensity below that for EV)—black letters on white background. Where appropriate, complexes are subdivided into core and putative members, and/or with canonical vs. non-canonical designations.

### Identification of LSD1-dependent GFI1B binding partners

LSD1 binding is necessary for GFI1B function (10, 43). However, discovering elements from functionally distinct transcriptional regulatory complexes within the GFI1 B proximitome suggests an important interplay between these elements and LSD1. Moreover, at least three of these complexes (BHC, NuRD and CtBP) contain LSD1 as a core component (58, 60, 80). Given the importance of LSD1 for GFI1B function, we hypothesized one or more transcriptional regulatory complexes represented in the GFI1B proximitome would be recruited in an LSD1-dependent manner, and if its components were depleted, would impair GFI1B function even when GFI1B—LSD1 binding potential is preserved. To address this hypothesis, we engineered expression constructs encoding fusion proteins comprised of BirA* and either LSD1 binding-competent forms of GFI1B (GFI1B-BirA*:HA and SNAG-BirA*:HA) or GFI1B variants with impaired LSD1 binding (GFI1B-P2A-BirA*:HA and GFI1B-ΔSNAG-BirA*:HA). A BirA*:HA construct served as an additional negative control (Fig. 5A). Fusion proteins were inducibly expressed in K562 cells and their biotinylating ability compared in whole cell lysates by SDS-PAGE, transblotting and SAv:HRP detection (Fig. 5B). Equivalent expression and robust, proteome-wide biotinylation were observed for each construct induced with doxycycline. We then tested the ability of each fusion protein to establish proximity relationships with known SNAG domain binding proteins, LSD1 and RCOR1 (43, 45). As expected, GFI1B-BirA*:HA and SNAG-BirA*:HA display clear enrichment for biotinylated LSD1 and RCOR1 in SAv-Sepharose pull downs, while LSD1 non-binding variants (P2A and ΔSNAG) show biotinylation comparable to background seen for the BirA*:HA negative control (Fig. 5C). Equally important, we observed BirA*:HA biotinylation comparable to each of its fusion proteins, suggesting equal capacity for spontaneous biotinylation and mitigating concerns that the fusion partner might adversely impact BirA* enzyme activity. Collectively, these results support use of GFI1B variants to unambiguously define the LSD1-dependent elements of GFI1 B complexes governing transcriptional control and cell fate specification.

**Figure 5.**
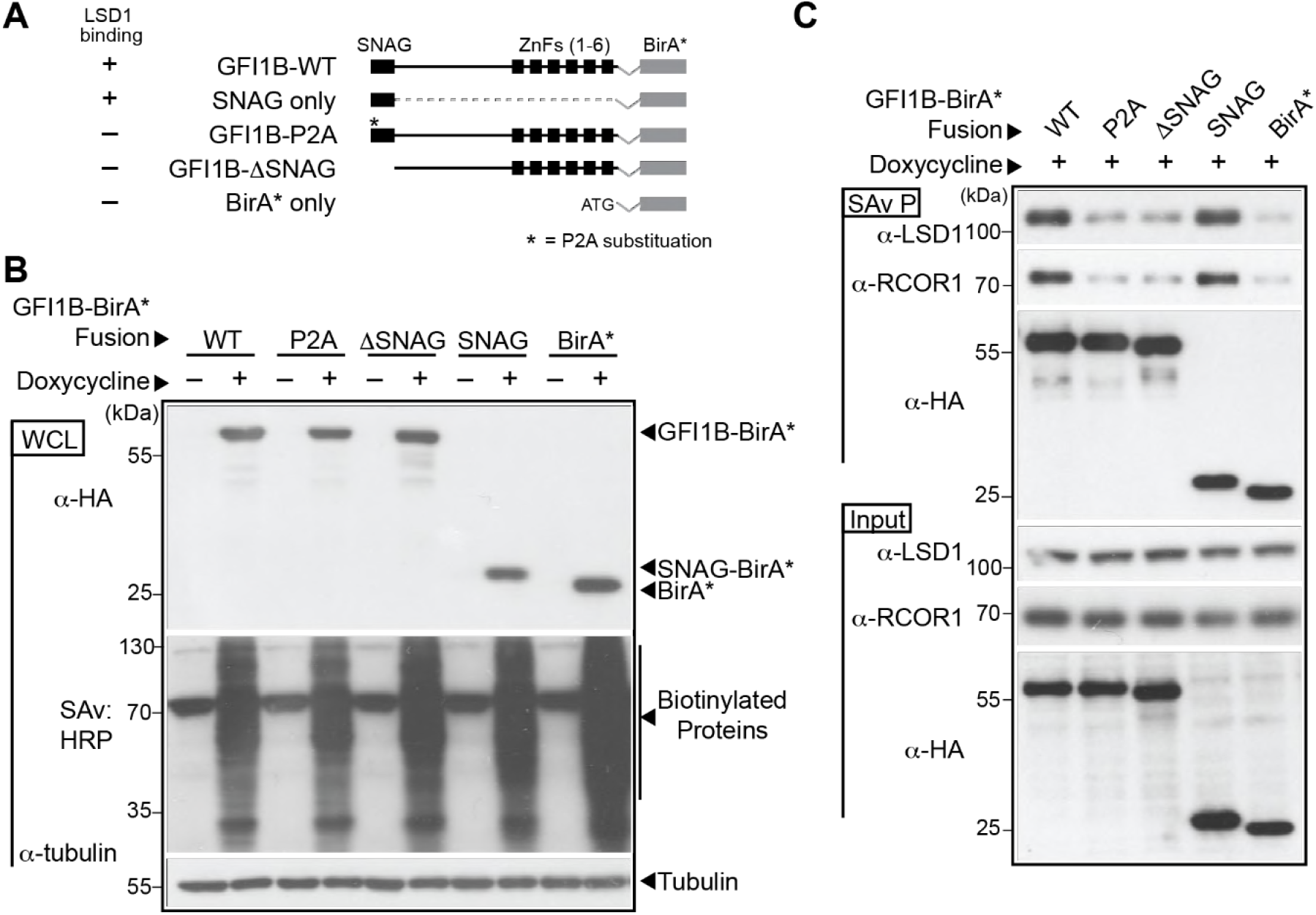
LSD1 non-binding variants of GFI1B-BirA* fail to biotinylate LSD1 and RCOR1. (A) Graphical representation of GFI1B-BirA*:HA fusion proteins employed in experiments. Each fusion protein is comprised of the GFI1B regions shown in frame with the BirA* expression cassette and a C-terminal HA epitope tag to insure equivalent expression of all forms. The BirA*:HA construct lacks GFI1B structures and instead begins with an ATG. Wild type (WT) GFI1B is shown. The SNAG and zinc finger (ZnF) domains (1–6) of GFI1B are shown as black boxes. A thin line represents the GFI1B linker and connections between ZnFs. The proline to alanine substitution (P2A) in the GFI1B SNAG domain is indicated with an *. ΔSNAG represents deletion of the SNAG domain but preservation of the remaining GFI1B primary structure. (B) Expression of GFI1B-BirA*:HA forms and proteome-wide biotin modification in K562 cells. K562 cells were transduced with lentivirus to inducibly express GFI1B-BirA*:HA forms shown and stable isolates selected as polyclonal populations. Doxycycline-inducible expression of each fusion protein was confirmed by western blot against the common HA epitope tag and biotinylating activity confirmed proteome-wide by SAv:HRP detection of transblotted total cellular protein. Tubulin served as a loading control. (C) LSD1 and RCOR1 are enriched among biotinylated proteins generated by BirA*:HA fusions competent for LSD1 binding. Biotin-modified proteins from K562 cells transduced with BirA*:HA fusion proteins shown were purified from whole cell extracts (SAv P), fractionated by SDS-PAGE and subjected to western blot with α-LSD1, α-RCOR1 and α-HA antibodies. Equivalent expression of each protein in whole cell lysates (Input) was confirmed by western blot with these same antibodies.

In pursuit of this goal, we deployed these tools for quantitative proximitome analyses to compare partnerships between GFI1B forms which are either capable of (GFI1B-BirA*:HA and SNAG-BirA*:HA), or deficient in (GFI1B-P2A-BirA*:HA, GFI1B-ΔSNAG-BirA*:HA and BirA*:HA) LSD1 binding. Fusion protein expression was induced by addition of doxycycline and biotinylated proteins collected on SAv-Sepharose beads as above. After high stringency washing, affinity purified proteins were analyzed by LC-MS-MS as described above. Triplicate read intensities were used to calculate an average sum read intensity for each protein identified in the mass spectral screen. Proteins were ranked according to average sum read intensities in the LSD1 non-binding control and fold enrichment for each protein was calculated as a ratio of average sum read intensities between LSD1 binding and non-binding variants. P-values were calculated from variation among the twenty nearest ranked neighbors for each protein in the data set. Compared to LSD1 non-binding variants GFI1B-ΔSNAG-BirA*:HA (Fig. 6A) and GFI1B-P2A-BirA*:HA (Fig. 6B), we observed enrichment for nine of the ten core (LSD1, RCOR1, PHF21A, HMG20B, HDAC1and HDAC2) and putative (GSE1, HMG20A, and ZMYM2) components of the BHC complex in the GFI1B-BirA*:HA proximitome. These results indicate BHC complex recruitment is lost with impaired GFI1B—LSD1 binding. To focus specifically on the SNAG—LSD1 interaction as a determinant of BHC complex recruitment, we determined enrichment for BHC complex components in the SNAG-BirA*:HA proximitome relative to BirA*:HA only. Again, we find enrichment for these nine BHC complex components in the SNAG-BirA*:HA proximitome.

**Figure 6.**
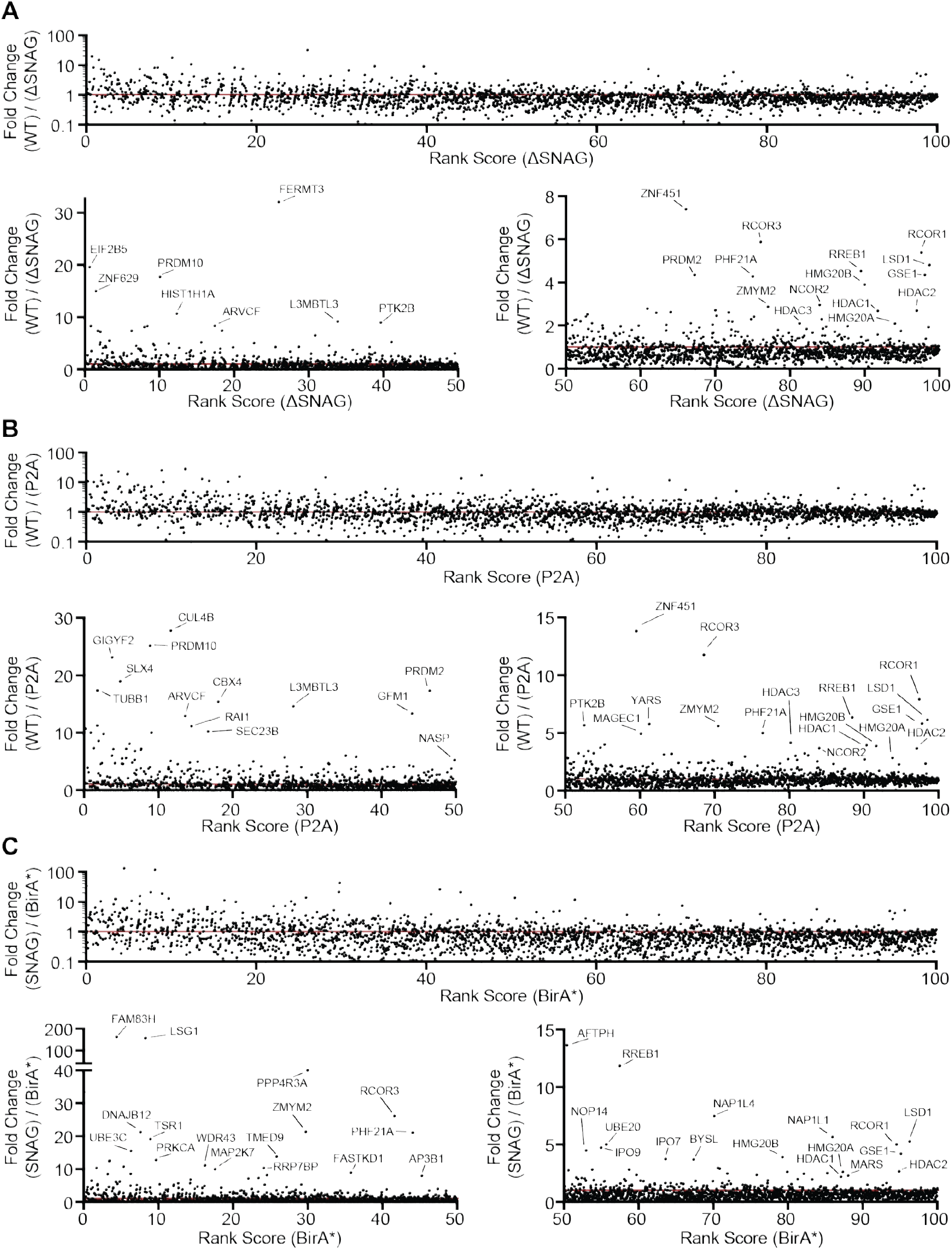
GFI1B recruits the BHC complex in an LSD1-depend manner via the SNAG domain. K562 cells stably and inducibly expressing the BirA*:HA fusion proteins shown in 5A were deployed to define GFI1B proximity partners whose recruitment is LSD1-dependent. Fusion protein expression was induced by doxycycline addition and biotinylation permitted *in situ*. Biotin-modified proteins were captured on SAv-Sepharose and analyzed by LC-MS-MS as described in Materials and Methods. (A-C) Dot plots are shown comparing fold change in average sum read intensity among fusion proteins with intact SNAG domains and capable of LSD1-binding (GFI1B-BirA*:HA and SNAG-BirA*:HA) to those with disrupted or absent SNAG domains and deficient in LSD1 binding (GFI1B-P2A-BirA*:HA, GFI1B-ΔSNAG-BirA*:HA, and BirA*:HA). For each comparison (A—GFI1 B-BirA*:HA vs. GFI1B-ΔSNAG-BirA*:HA, B—GFI1 B-BirA* vs. GFI1B-P2A-BirA*:HA, and C—SNAG-BirA*:HA vs. BirA*:HA), average sum read intensities were ranked according to the LSD1-nonbinding sample and plotted on the x-axis. Fold change in average sum read intensities, represented by a ratio between the LSD1-binding and LSD1-nonbinding sample in each comparison, is plotted on the y-axis. Each comparison, A-C, is represented by three panels. On top in each is the entire data set for that comparison, and below are two panels dividing the percentile rank score from 0-50 and from 50-100 with expanded y-axis limits to facilitate visualization of proteins enriched in the data set for the LSD1-binding form. Outlying proteins are labeled with their gene names. A red line indicates equal read intensities, and thus no enrichment, for the LSD1-binding over the LSD1-nonbinding sample.

LSD1, HDAC1, and HDAC2 are shared among the BHC, CtBP and NuRD complexes, while RCOR1 is found in both BHC and CtBP complexes. To confirm the LSD1-dependence of BHC complex recruitment specifically, we surveyed the top 100 enriched proteins in each binary proximitome comparison (GFI1B-BirA*:HA vs. GFI1B-ΔSNAG-BirA*:HA, GFI1B-BirA*:HA vs. GFI1 B-P2A-BirA*:HA, and SNAG-BirA*:HA vs. BirA*:HA) for the unique components of BHC, CtBP and NuRD complexes (Table 1 and Table S1). While unique components of the BHC complex were found in each binary proximitome comparison performed, none of those unique to the CtBP or NuRD complexes were observed. These findings indicate that among LSD1-containing complexes detected in the GFI1B proximitome, only the BHC complex displays LSD1-dependence.

**Table 1.**
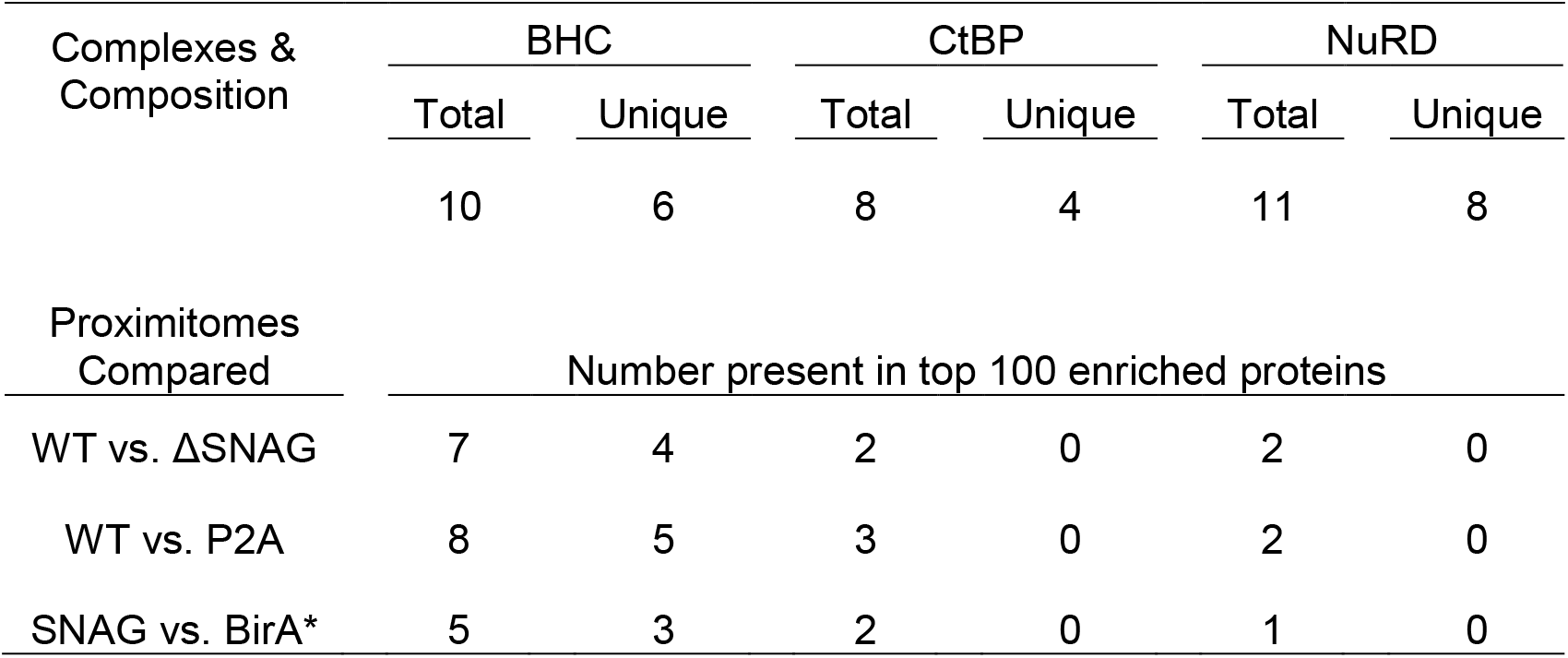
LSD1-dependent components of GFI1B proximitomes. Shown are components of LSD1-containing transcription complexes represented in the GFI1B proximitome. The core and unique components of each complex are enumerated (top), and the number of core and unique components from these complexes found in each proximitome comparison is shown.

| Complexes & Composition | BHC |  | CtBP |  | NuRD |  |
| --- | --- | --- | --- | --- | --- | --- |
|  | Total | Unique | Total | Unique | Total | Unique |
|  | 10 | 6 | 8 | 4 | 11 | 8 |
| Proximitomes Compared | Number present in top 100 enriched proteins |  |  |  |  |  |
| WT vs. ΔSNAG | 7 | 4 | 2 | 0 | 2 | 0 |
| WT vs. P2A | 8 | 5 | 3 | 0 | 2 | 0 |
| SNAG vs. BirA* | 5 | 3 | 2 | 0 | 1 | 0 |

### GFI1B-mediated transcriptional repression and cell fate require the BHC complex

To confirm findings from mass spectral screens, we first assessed associations between GFI1B and candidate BHC complex components and tested their dependence upon GFI1B—LSD1 binding. We focused on unique elements of the BHC complex. GFI1B-BirA*:HA, GFI1B-P2A-BirA*:HA, GFI1B-ΔSNAG-BirA*:HA or SNAG-BirA*:HA were inducibly expressed in K562 cells and biotinylated proteins purified on SAv-Sepharose beads. HMG20A, HMG20B, and PHF21A were then quantified among purified proteins by western blot. Each was enriched among proteins purified from cells expressing GFI1B-BirA* fusions capable of LSD1 binding relative to non-binding controls (Fig. 7A). Notably, no such enrichment was observed for SMARCB1 or SMARCC1, members of the SWI/SNF complex that showed enrichment but no LSD1 dependence in GFI1B proximitome analyses. Nor did SMARCB1 or SMARCC1 show a proximity relationship when SNAG-BirA*:HA was employed, confirming a distinct and LSD1-independent means of associating with GFI1B. Using flag-tagged wild type and -P2A forms of GFI1B, we extended findings from proximity labeling studies to traditional coprecipitation assays. For HMG20A, HMG20B, and PHF21A, binding to GFI1B-P2A was significantly impaired relative to wild type GFI1B, confirming dependence upon LSD1 recruitment (Fig. 7B-D). This observation is further supported by co-precipitation assays showing interactions with LSD1 for both HMG20A and HMG20B (Fig. 7E).

**Figure 7.**
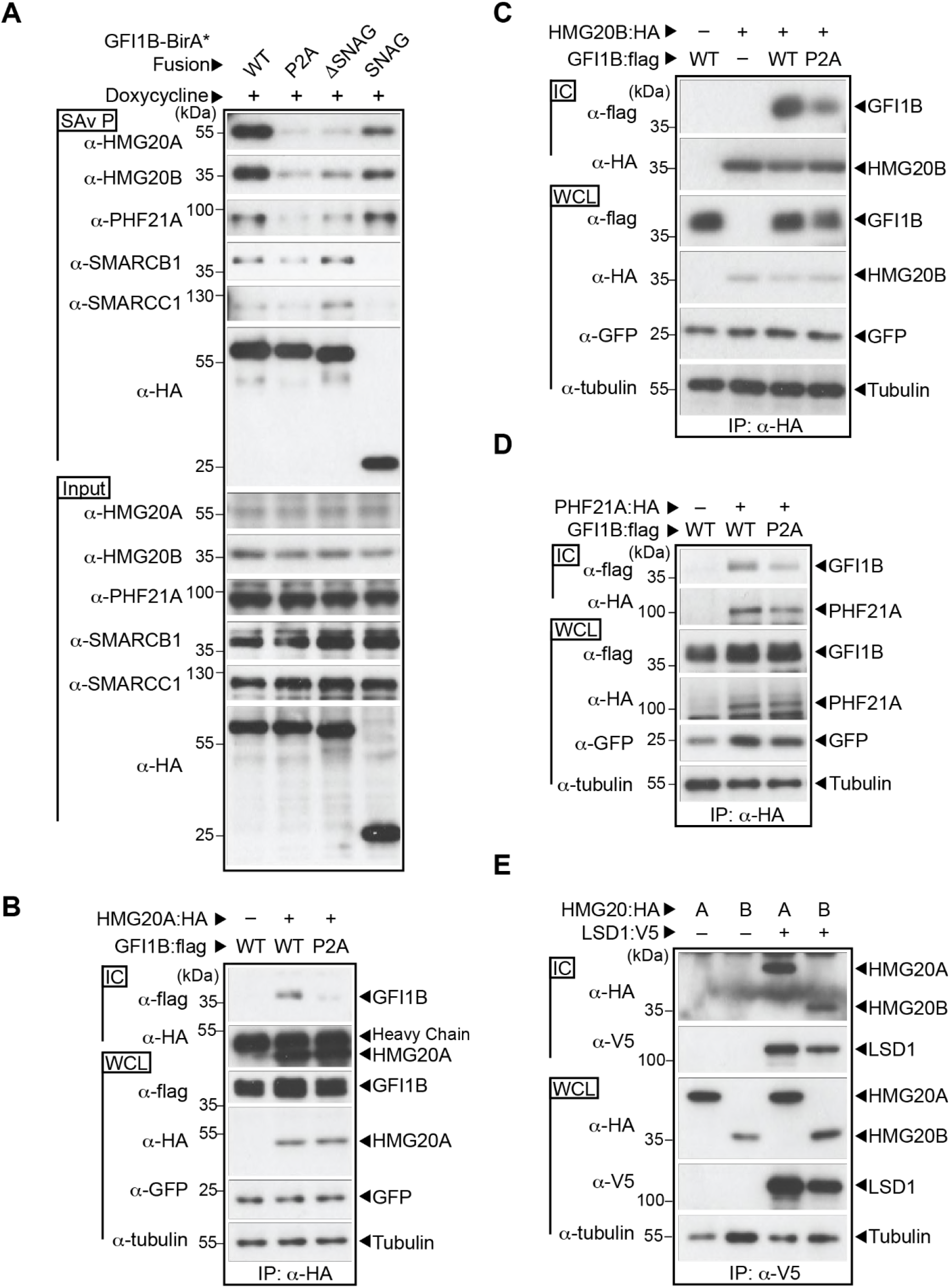
BHC complex components HMG20A, HMG20B and PHF21A bind GFI1 B in an LSD1-dependent manner. (A) BHC complex components are preferentially enriched in proximity labeling assays with fusion proteins competent for LSD1 binding. Biotin-modified proteins were purified from K562 cells inducibly expressing the BirA*:HA fusion proteins shown. HMG20A, HMG20B, PHF21A, SMARCB1 and SMARCC1, as well as the BirA*:HA fusion proteins, were quantified in SAv-Sepharose pulldowns (SAv P) by western blot. Equivalent input for each protein was confirmed by western blot of whole cell lysates (Input). (B-D) GFI1B binding to HMG20A, HMG20B and PHF21A require GFI1B—LSD1 binding. COS7L cells were transfected with HA-tagged HMG20A (B), HMG20B (C), or PHF21A (D) and wild type (WT) or -P2A variants of GFI1 B:flag. HMG20 and PHF21A proteins and their binding partners were purified in α-HA immune complexes. Co-precipitating GFI1B forms were detected in immune complexes by western blot using α-flag antibody. Equivalent expression and precipitation were evaluated in WCLs and ICs, respectively. GFP served as a transfection control. (E) HMG20A and HMG20B bind LSD1. COS7L cells were transfected with HA-tagged HMG20A or HMG20B along with a V5-tagged LSD1. LSD1 and its binding partners were collected in α-V5 immune complexes (IC). Co-precipitating HMG20A and HMG20B were detected in immune complexes by western blot using α-HA antibody. Equivalent expression and precipitation were evaluated in the whole cell lysates (WCL) and ICs, respectively.

LSD1 is required for both GFI1B function and BHC complex recruitment, but whether LSD1 exerts its impact through BHC complex recruitment is not known. To address this question, we targeted unique BHC complex components for depletion in K562 cells then tested GFI1B’s ability to direct changes in cell fate and gene expression (Fig. 8). HMG20A and HMG20B are known to bind structured elements in DNA in a sequence non-specific manner (81–83). They have been shown to have an oppositional role in neuronal differentiation via modulation of REST-responsive genes (80, 84). Their contributions to hematopoietic differentiation have not been described. HMG20A and HMG20B were depleted alone or together using inducible shRNA constructs (Fig. 8A), then effects of enforced GFI1B expression on K562 cell differentiation and gene expression were assessed. While neither HMG20A nor HMG20B depletion alone had a notable impact on K562 cell hemoglobinization, simultaneous depletion of both factors prevented this GFI1B-driven phenotype (Fig. 8B). Likewise, impaired TPA-induced megakaryocytic differentiation brought on by enforced GFI1B expression was not significantly affected by depletion of either HMG20A or HMG20B alone. Yet, their concurrent depletion enabled accumulation of CD61 on the K562 cell surface despite GFI1B overexpression (Fig. 8C). Erythroid differentiation blockade brought on by concurrent depletion of HMG20A and HMG20B correlated with reduced expression of genes required for heme and globin chain synthesis and with loss of repression of GFI1 B target genes (Fig. 8D-F). Notably, LSD1 expression is unaffected by HMG20A or HMG20B depletion (Fig. 8A). Thus, HMG20A and HMG20B make functional contributions to LSD1-dependent outcomes driven by GFI1B. Their concurrent absence impairs GFI1B-driven erythroid fate in K562 cells and enables alternate, TPA-driven megakaryocytic differentiation in the context of enforced GFI1B expression that would normally block it. Moreover, unlike the oppositional relationship between HMG20A and HMG20B observed in REST-regulated neural differentiation, these findings intimate functional redundancy between them in GFI1 B-mediated transcription and its control of cellular identity.

**Figure 8.**
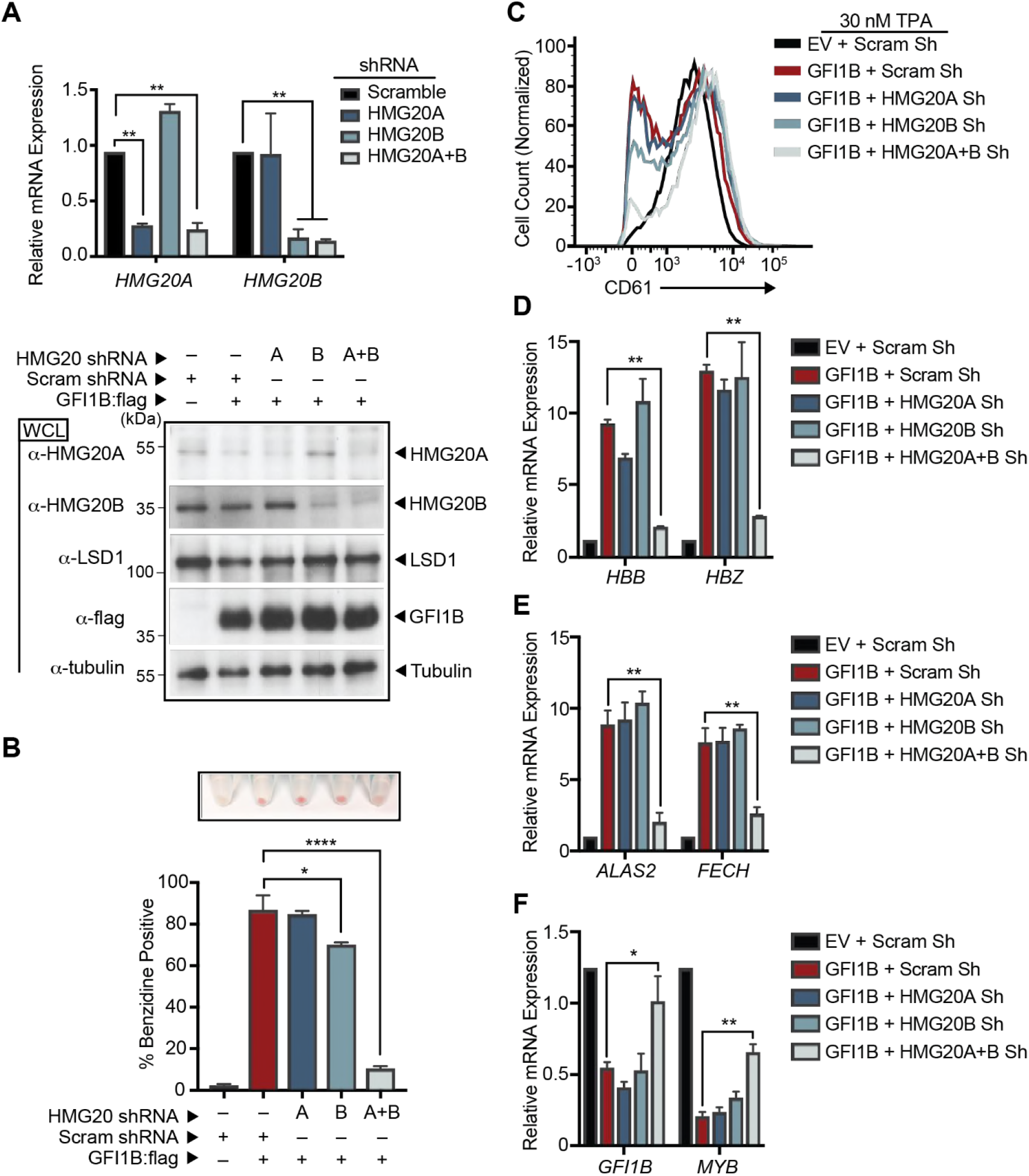
HMG20A and HMG20B are required for GFI1B-mediated cell fate changes in K562 cells. (A) HMG20A and HMG20B depletion in K562 cells. RNA (Top) and Protein (Bottom) were collected from K562 cells transduced with shRNA targeting HMG20A, HMG20B, HMG20A and HMG20B (HMG20A+B), or a content-matched scrambled control. HMG20A and HMG20B expression levels were evaluated by RT-qPCR and normalized to *GUS.* HMG20A, HMG20B, LSD1, and GFI1B:flag levels were determined by western blot. Tubulin served as a loading control. (B-F) HMG20A/B depletion abolishes GFI1B-dependent cell fate determination in K562 cells. K562 cells were transduced with inducible shRNAs targeting HMG20A, HMG20B, HMG20A and B or a scrambled control. The shRNAs were induced with doxycycline, cells were infected with GFI1 B:flag-expressing retrovirus and erythroid differentiation allowed to proceed. (B) Benzidine staining was quantified by counting positive cells in bright field microscopy while hemoglobinized cell pellets are shown above each condition shown in the bar graph. (C) GFI1 B-dependent inhibition of TPA-induced CD61 expression requires HMG20A/B. K562 cells overexpressing GFI1B and depleted of HMG20A, HMG20B, or HMG20A and B were treated with TPA and cell surface expression of CD61 quantified by flow cytometry. Histograms presented are representative of three independent experiments. (D-F) mRNA for globin genes *HBB* and *HBZ* (D), heme biosynthesis genes *ALAS2* and *FECH* (E) and GFI1B-repressed target genes *GFI1B* and *MYB* (F) were quantified by RT-qPCR normalized to *GUS* reference gene. Results are expressed as mean ± 2SD. Statistical significance was determined by two-sided unpaired *t*-test; *, *p* < 0.05; **, *p* < 0.005; ****, *p* < 0.00005.

To extend observations made with HMG20A and HMG20B, we tested the effects of GSE1 depletion on GFI1B functions in transcriptional repression and erythroid fate specification in K562 cells. GSE1 is a proline-rich coiled-coil domain protein implicated in pathogenesis of breast and gastric cancers (85, 86). It is widely expressed in human cells, but its role in cell fate decisions in hematopoiesis have not been studied. As for HMG20 proteins, reduced GSE1 expression blocks erythroid differentiation and gene expression changes in K562 cells which depend upon the GFI1B—LSD1 binding relationship (Fig. 9). Together with results from LSD1, HMG20A and HMG20B depletion, these data highlight that LSD1 is necessary but not sufficient for GFI1B-driven outcomes and showcase a critical role for the BHC complex as an LSD1-dependent determinant of GFI1B-mediated transcriptional repression and cell fate decisions in hematopoiesis.

**Figure 9.**
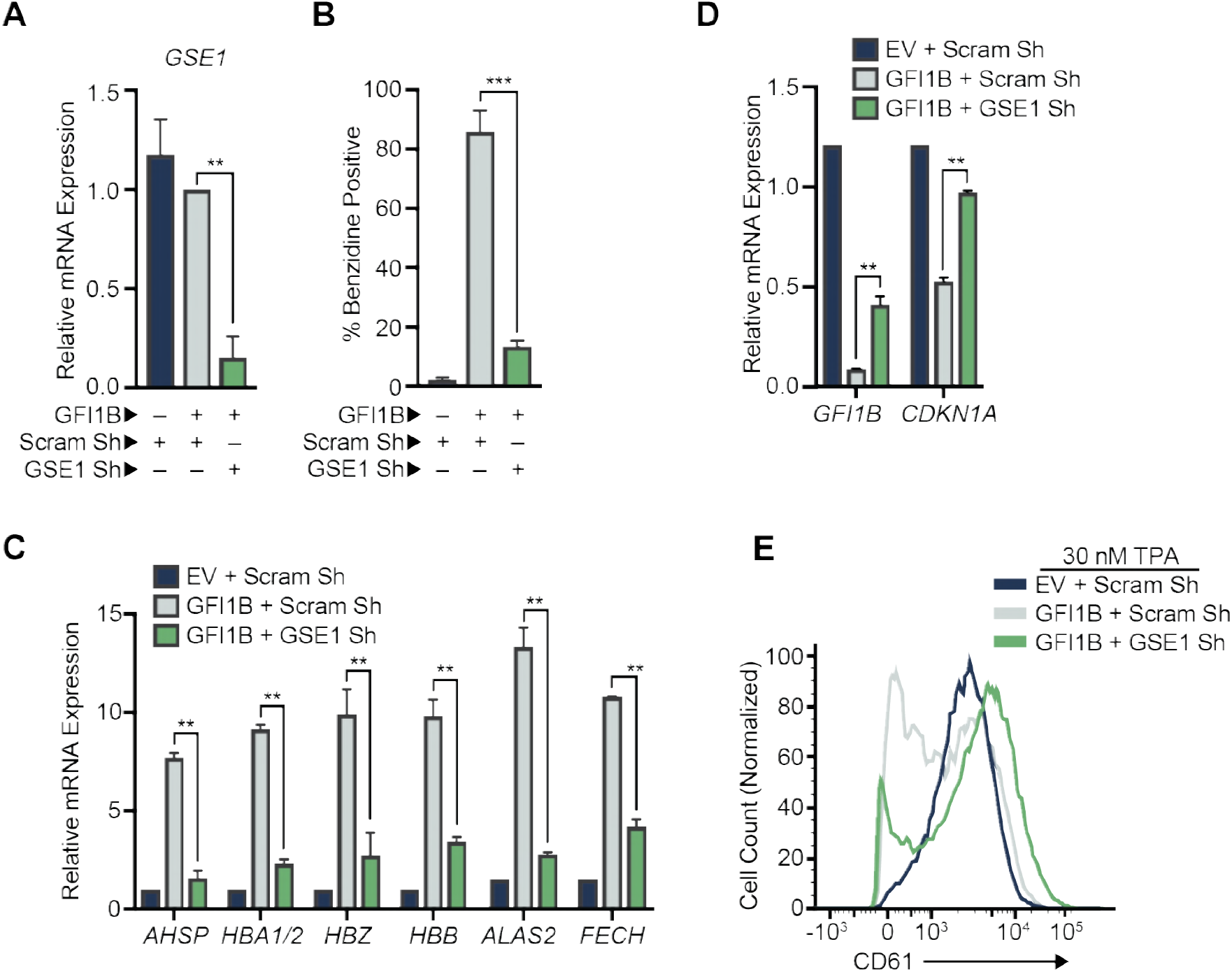
GSE1 is required for GFI1B-mediated cell fate changes in K562 cells. (A) GSE1 depletion in K562 cells. RNA was collected from K562 cells transduced with shRNA targeting GSE1 or a content-matched scrambled control. GSE1 expression levels were evaluated by RT-qPCR and normalized to *GUS.* (B-D) GSE1 depletion abolishes GFI1B-dependent cell fate determination in K562 cells. K562 cells were transduced with inducible shRNAs targeting GSE1 or a scrambled control. The shRNAs were induced with doxycycline, cells infected with GFI1B:flag expressing retrovirus and erythroid differentiation allowed to proceed. (B) Benzidine staining was quantified by counting positive cells in bright field microscopy. (C-D) mRNA for globin and related genes *AHSP, HBA1/2, HBZ*, and *HBB;* heme biosynthesis genes *ALAS2* and *FECH* (C); and GFI1B-repressed target genes *GFI1B* and *CDKN1A* (D) were quantified by RT-qPCR and normalized to *GUS* reference gene. Results are expressed as mean ± 2SD. (E) GFI1B-dependent inhibition of TPA-induced CD61 expression requires GSE1. K562 cells overexpressing GFI1B and depleted of GSE1 were treated with TPA and cell surface expression of CD61 quantified by flow cytometry compared to control cells. Histograms presented are representative of three independent experiments. Statistical significance was determined by two-sided unpaired *t*-test; **, *p* < 0.005; ****, *p* < 0.00005.

## DISCUSSION

The control of gene expression involves DNA binding proteins that recognize discrete sequence elements in regulatory regions of genes, directing site-selective assembly of multiprotein complexes that modulate chromatin structure, and ultimately, access to coding regions by the basal transcription machinery (87, 88). By deploying individual elements in a modular fashion, cells insure flexibility and economy as they orchestrate transcriptional programs to control the specification of cell fate. Proper function of these complexes is layered upon a spatial and temporal order of events that combine to create a flow of information needed for a specific outcome among the myriad of possibilities. Yet, this strategy necessarily creates dependency relationships between factors cooperating to accomplish functions required for an outcome to be realized. Malfunction of just one complex component, or disordered operation of a complex in space or time may impair functional outputs and thus mask the underlying molecular complexities required to achieve them.

The relationship between GFI family proteins and LSD1 exemplifies this theme. Both GFI1 and GFI1B require LSD1 binding for transcriptional repression and point mutations in their SNAG domains completely abolish activity in multiple functional assays (43, 45, 46, 89). It is not, however, immediately apparent that other factors might enable LSD1’s central role, and identifying these collaborators is technically challenging. We deployed an unbiased, proteome-wide proximity labeling strategy based upon a promiscuous variant of BirA (BirA*) to prospectively and systematically label with biotin factors that dwell in the vicinity of GFI1B and then to determine functionally relevant partners which are recruited in a manner requiring LSD1, its obligate effector. In so doing, we show that while LSD1 recruitment is critical to GFI1B-driven erythroid differentiation, it does not operate alone as its singular depletion in K562 cells may suggest. Rather, LSD1 is responsible for selective recruitment of the BHC complex, implying that its contributions to GFI1B-mediated outcomes may reflect both its intrinsic demethylase activity and its role as a scaffolding component for this multiprotein complex. A model summarizing the collaborative relationship between LSD1, BHC complex components and other partners in GFI1/1B-mediated transcriptional repression is shown (Fig. 10).

**Figure 10.**
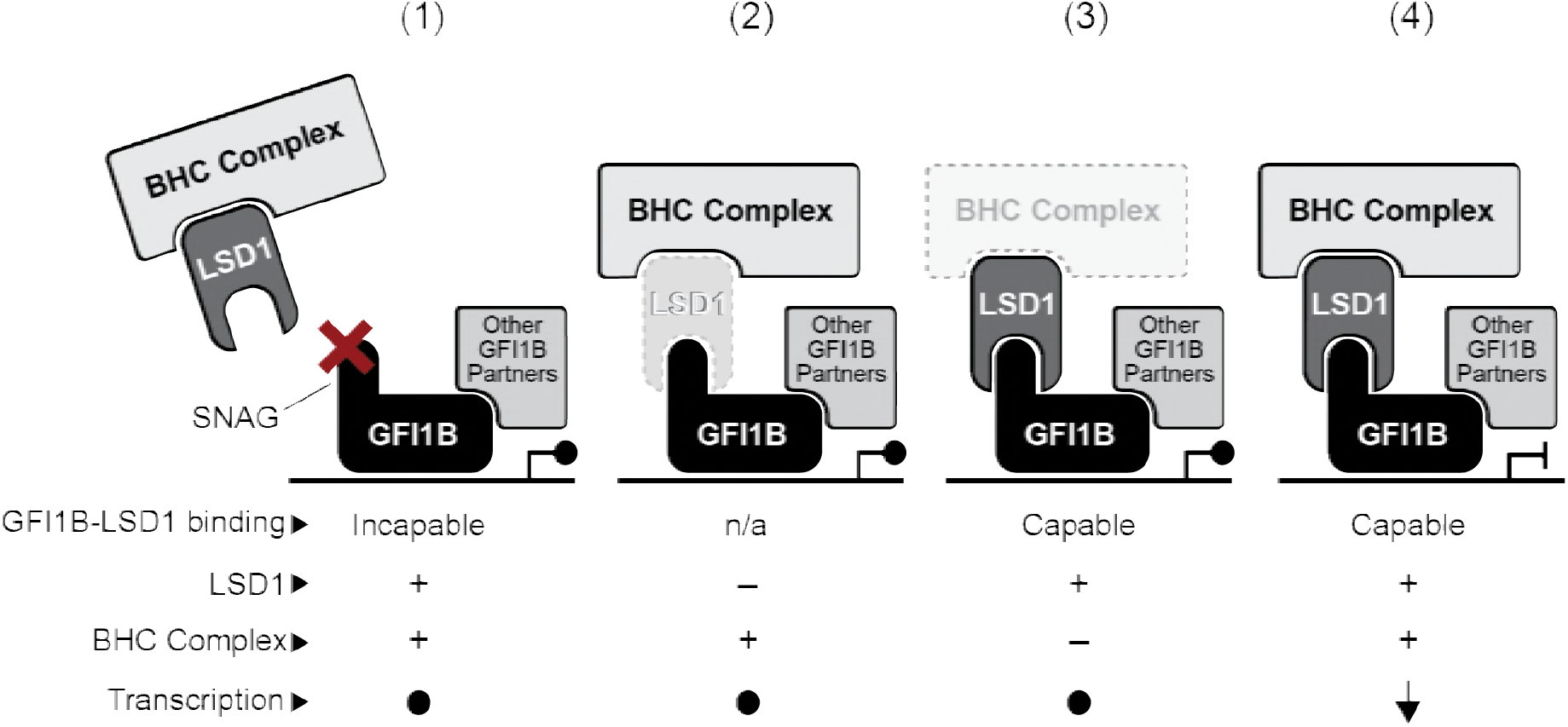
A working model for contributions of the BHC complex to GF1B-mediated repression. LSD1 serves as a bridge between GFI1B and other components of the BHC complex, including HDAC1/2, HMG20A, HMG20B, PHF21A, RCOR1, ZMYM2, ZNF217, and GSE1. Therefore, BHC complex recruitment is rendered LSD1 dependent. A functional GFI1B transcriptional repression complex requires coincident recruitment of these factors, along with other partners brought to the promoter in an LSD1-independent manner by GFI1B or DNA binding proteins with which it collaborates. Dysfunction resulting from (1) an impaired, SNAG-dependent GFI1B—LSD1 recruitment mechanism (2) LSD1 depletion or intrinsic change that disrupts SNAG binding or (3) ineffective formation or operation of BHC complex components could each poison the GFI1B—LSD1 functional axis to create a permissive state for misexpression of GFI1B-regulated genes. Only when each element is present and functioning properly (4) is the appropriate repressive outcome achieved. Trans-regulation by each component toward the others, or toward partners that engage GFI1B in a LSD1-independent manner, may also make important contributions to enabling and fine-tuning target gene expression.

LSD1 displays *in vitro* demethylase activity toward peptides containing mono- and di-methylated lysines (K), best exemplified by demethylation of H3K4me1/me2, and toward purified histones (48, 90). However, in a chromatinized context, methylated histones are poor substrates for LSD1 (91). Efficient demethylation of chromatinized histones by LSD1 requires concurrent RCOR1 binding, and this activity can be further stimulated by an interaction between RCOR1 and SUMO2/3 (91, 92). Likewise, LSD1’s demethylase activity is favored by HDAC1-mediated deacetylation of LSD1-K374ac (93). Observations such as these support the notion that LSD1’s demethylase activity may hinge upon its inclusion in multiprotein complexes to enable its proper post-translational modifications and allosteric modulation. Additionally, by serving as a scaffold, LSD1 may enable regulation of other components of the BHC complex, or perhaps discrete regulatory modules recruited to operationalize GFI1B activity as a transcriptional repressor. Absent these cooperative relationships, LSD1 may fail to execute its obligate effector role for GFI1B despite the means of its recruitment by GFI1B being intact. Investigating these possibilities presents a daunting challenge, but one that can now be tackled using the situational capabilities of proximity labeling approaches. Moreover, a holistic view of LSD1 contributions to GFI1B’s repressor and cell fate specifying activities is consistent with both its pivotal role and its dependence upon partnerships.

To establish evidence supporting this view, we focused on BHC complex components HMG20A and HMG20B as LSD1 partners. This focus was stimulated by studies in *Schizosaccharomyces pombe* concerning the role of Lsd1/2 in replication fork pausing associated with mating type class switching (94). Lsd1/2 is required for growth and viability in S. *pombe.* Notably, S. *pombe* strains rendered catalytically inactive at *lsd1/lsd2* are viable and do not phenocopy *lsd1/lsd2* deletion, suggesting they have a non-enzymatic role. Moreover, Lsd1 contains a C-terminal HMG domain that is lacking in mammalian LSD1, and HMG domain mutations impair mating type class switching through defective imprinting at the *mat1* locus brought on by dysfunctional replication fork pausing (48, 94). The importance of the Lsd1 HMG domain to a phenotype dependent upon structural change in DNA, the absence of the HMG domain in mammalian LSD1, the suggestion of a non-enzymatic role for Lsd1 and the inclusion of HMG20 proteins in the GFI1B proximitome made HMG20A and HMG20B attractive candidates in GFI1B—LSD1-dependent outputs in K562 cells. Our finding that concurrent HMG20A/B depletion blocks LSD1-dependent functions of GFI1B suggests that their inclusion in the BHC complex is essential to enable LSD1 actions. Similar observations made by depleting GSE1 reinforce this notion. Notably, HMG20A and HMG20B are ubiquitously expressed (95). They contain N-terminal HMG boxes which bind DNA in a sequence non-specific manner and C-terminal coiled-coil domains required for dimerization and incorporation into multiprotein complexes (67, 84). In neural development, HMG20A and HMG20B display an oppositional relationship at REST-regulated genes (65, 80, 84). Yet, our data suggest these proteins may also compensate for deficiencies in one another and that perhaps the relative abundance of each may fine tune transcriptional activity of the BHC complex at sites to which it has been recruited. Thus, with site-selective DNA binding proteins (e.g., GFI1B), epigenetic modifiers (e.g., LSD1, HDAC1/2), their cofactors (e.g., RCOR1), and readers of epigenetic state (e.g., PHF21A) as partners, a network of interdependent proteins may cooperate to direct the quality, quantity and timing of transcriptional programs that govern the specification of cell fate. Our findings provide a framework for exploring the interdependent and situational relationships among these classes of proteins.

## MATERIALS AND METHODS

### Cell culture and viral transduction

COS7L, HEK293T (American Type Culture Collection), MEL and 293-T-Rex-5xGal-luciferase cells (generously provided by Raphael Margueron) were maintained in Dulbecco’s modified Eagle’s medium (DMEM) supplemented with 10% fetal bovine serum (FBS). K562 cells were maintained in Roswell Park Memorial Institute (RPMI) medium supplemented with 10% FBS and 25 μM HEPES. All cell culture media was supplemented with 2 μM L-Glutamine, 50 units/ml penicillin, and 50 μg/ml streptomycin. All cell media and additives were obtained from Thermo-Fisher. All cell lines were verified using the Geneprint 10 system (Promega). For viral production, HEK293T cells were transfected with a viral vector containing the transgene along with the respective lenti-(psPAX2 and pMD2G) or retroviral (gag/pol and VSVG) packaging plasmids using polyethylenimine (PEI) (3 μg PEI:1 μg DNA). Viral supernatant was collected 48- and 72-hours posttransfection and clarified by filtration (Corning, 45 μm). K562 cells were transduced with viral particles in the presence of 8 μg/ml polybrene for 16 hours. Stable cells were obtained by selecting with 5 μg/ml puromycin for 4 days, 400 μg/ml G418/neomycin or hygromycin for 10 days. For LSD1 and HMG20B small hairpin (shRNA) experiments, K562 cells were transduced with Tet-pLKO-neo (a gift from Dmitri Wiederschain, Addgene plasmid #21916) containing either LSD1-, HMG20B-targeted shRNA or scrambled control and selected in neomycin to establish stable isolates. For HMG20A and GSE1 shRNA experiments, K562 cells were transduced with EZ-Tet-pLKO-Hygro (a gift from Cindy Miranti, Addgene plasmid #85972) containing either HMG20A-, GSE1-targeted shRNA or scrambled control and selected in hygromycin. To obtain double HMG20A and B shRNA knockdown, K562 cells were first transduced with either HMG20B or scrambled shRNA vector, selected with neomycin, and subsequently transduced with either HMG20A or scrambled shRNA vector and selected with hygromycin. shRNA expression was achieved by treating transduced cells with 500 ng/ml doxycycline every 24 hours. shRNA sequences are listed in Table S2.

### RNA extraction and gene expression analysis

Total RNA was prepared using RNeasy (Qiagen), as per the manufacturer’s instructions, including on-column DNAse I digestion to avoid potential gDNA contamination. The cDNA was generated from 100-300 ng of total RNA using iScript Reverse Transcription Supermix for RT-qPCR (Bio-Rad). cDNA was diluted 2X with water, and 1 μl of cDNA was used as a template for real-time PCR (qPCR). Quantitation was performed by qPCR with the CFX96 Real Time System (Bio-Rad), using 5 μl SsoAdvanced Universal SYBR Green Supermix (Bio-Rad), and 330 nM of the indicated forward and reverse primers in a 10 μl reaction. Relative mRNA expression was determined by the ΔΔCt method and normalized to a *GUS* reference gene. Each experiment was performed in triplicate. Statistical significance was determined by two-sided unpaired *t*-tests using GraphPad Prism 7.0. Sequences of all oligonucleotides used are listed in Table S2.

### Constructs and subcloning

GFI1B (NM_008114.3), PHF21A (NM_008114.3), HMG20A (NM_025812.3), and HMG20B (NM_010440.3) were amplified by PCR (Platinum SuperFi DNA Polymerase, Invitrogen) from cDNA obtained from MEL cells and subcloned into pcDNA3.1(-) (Invitrogen) with either a 3x-flag, HA, or V5 tag on the 3’ end of the gene. The GFI1B-P2A mutation was made by site-directed mutagenesis. Retroviral GFI1B:3x-flag vectors were produced by subcloning GFI1B into the pMSCV (murine stem cell virus)-IRES-Puro vector using EcoR1. GFI1B fusion proteins to Gal4 were generated by PCR amplification of the GFI1B SNAG domain (amino acid 1-20) or GFI1B-Δ1C (amino acids 1-244 only— eliminating ZnF 4, 5 and 6, which remove intrinsic DNA binding via GFI1B sequence) and inserted into pcDNA3.1 N-terminally and in frame with the Gal4 DNA-binding domain (amino acids 1-147) by EcoR1/Xba1. GFI1B fusion proteins with BirA* were made by subcloning GFI1B-WT, -P2A, -ΔSNAG (amino acids 21-330), or the SNAG domain (amino acids 1-20) into EcoR1/BamH1 sites in frame to BirA* in the MCS-BioID2-HA vector, a gift from Kyle Roux (Addgene plasmid #74224). Subsequently, GFI1B-BirA*:HA forms were expressed from PCR amplified sequences subcloned into pLVX-Tight-Puro (Clontech) with Not1/Mlu1 for doxycycline inducible expression with the Tet-On Advanced (Clontech) system. BirA*:HA only vector was created by inserting a start codon and an EcoR1 site into Age1/BamH1 (5’ of BirA*:HA) in pcDNA3.1 and subsequently put into pLVX-Tight-Puro by EcoR1. Primers used for cloning are listed in Table S2. The LSD1 construct was a generous gift from Sunil Sharma. Complete open-reading frame sequences for all constructs were confirmed by automated di-deoxy sequencing in the University of Utah DNA sequencing HSC Core Facility.

### Reagents and antibodies

Mouse monoclonal α-Flag (M2) and α-tubulin (clone B-5-1-2) were obtained from Sigma. Rabbit polyclonal α-HA (ab9110) and α-HMG20B (ab167415) were obtained from Abcam. Rabbit polyclonal α-LSD1 (C69G12) and α-RCOR1 (D6I2U) were obtained from Cell Signaling Technology. Mouse monoclonal α-HMG20A (D-5) and α-GFP (B-2) were obtained from Santa Cruz Biotechnology. Rabbit polyclonal α-SMARCB1 (A301-087A) and α-PHF21A (A303-603A) were obtained from Bethyl Laboratories and α-SMARCC1 (17722-1-AP) was obtained from Proteintech. Mouse monoclonal α-V5 (46–0705) was obtained from Invitrogen. Horse Radish Peroxidase (HRP) conjugated α-mouse (715-035-150) and α-rabbit (711-035-152) antibodies were obtained from Jackson Immunoresearch. HRP-conjugated streptavidin (RPN1231) (SAv) was obtained from GE Healthcare UK Limited. APC α-human CD61 (VI-PI.2), and 4’,6-diamidino-2-phenylindole, dilactate (DAPI) (422801) were obtained from BioLegend. Restriction endonucleases and ligases were obtained from New England Biosciences. Puromycin (puro), hygromycin (hygro), and G418/Geneticin (Neo) were obtained from Invitrogen. Protein-G Sepharose 4 Fast Flow and SAv-Sepharose beads were obtained from GE Healthcare Bio-Sciences AB. Biotin was obtained from TCI Chemicals and doxycycline (doxy) was obtained from Fisher Scientific. Benzidine di-hydrochloride, 12-*O*-Tetradecanoylphorbol-13-acetate (TPA), and polybrene were obtained Sigma. Polyethylenimine (PEI) 25kD Linear Form was obtained from Polysciences (#23966-2). All other materials were of reagent grade.

### Transient transfection and co-immunoprecipitation

Transient transfections in COS7L cells were performed using expression plasmids in the indicated combination and Lipofectamine 2000 per the manufacturer’s instructions. Monolayers were washed twice with phosphate-buffered saline (PBS), scraped into ice-cold lysis buffer (50 mM Tris, 150 mM NaCl, 10% glycerol, 1 mM EDTA, 1 mM dithiothreitol [DTT], 1% Triton X-100, 0.1% SDS, 1 mM phenylmethylsulfonyl fluoride [PMSF], 10 μg/ml aprotinin [pH 8.0]), disrupted by sonication (10 1-second pulses at 5W using Virtis Virsonic sonicator), rocked 30 min at 4°C, and clarified by centrifugation. As indicated by the figures, antibodies were added to clarified extracts for 90 min, then the immune complexes were captured by Protein-G-Sepharose beads that were washed in lysis buffer. Immune complexes were washed 4X with lysis buffer and boiled in SDS-PAGE sample buffer supplemented with DTT.

### Immunoblotting

After separation by SDS-PAGE, cleared lysates or immune complexes were examined by immunoblot following standard methods. Briefly, separated protein products were transferred to nitrocellulose filters (0.45 μm, Genesee Scientific) using 25 mM Tris, 192 mM glycine (pH 8.35) for 1 hour at 4°C. Nitrocellulose filters were blocked with PBS-T (1X PBS, 0.05% Tween-20) supplemented with 0.25% gelatin and 0.2% NaN3 (GBB) and incubated with primary antibodies overnight at 4°C as indicated in the figures. Filters were then washed successively with PBS-T and incubated with species-appropriate HRP-conjugated antibody for 1 hour at room temperature. After PBS-T washes, proteins were visualized by chemiluminescence detection (ECL; 30 mM Tris [pH 8.5], 375 μM Luminol, 67.5 μM p-Coumaric acid, 0.02% H2O2) using autoradiography film (Genesee Scientific).

### Luciferase reporter assays

Luciferase assays were performed using the Dual-Luciferase Reporter Assay System (Promega) as per manufacturer’s instructions. Briefly, 3.25 × 10^5^ 293-T-Rex-5xGal-luciferase cells were plated per well in 6-well plates. Transient transfection of the respective GFI1B:Gal4 expression constructs was performed with Lipofectamine 2000 as per manufacturer’s instructions. Reporter activity was measured in cell lysates collected 48 hours post transfection using a Modulus Microplate reader (Turner BioSystems). Firefly luciferase activity was normalized to constitutively expressed, co-transfected *Renilla* luciferase control. Statistical significance was determined by two-sided unpaired *t*-tests using GraphPad Prism 7.0.

### Chemical differentiation and phenotypic assays

4×10^5^ K562 cells were treated with 30 nM TPA. Cells were harvested 4 days postinduction, washed and incubated with α-CD61 antibody diluted 1:100 in Hanks Buffered Saline Solution (HBSS, Gibco) supplemented with 0.5% BSA (RPI Corp). Fluorescence signal from DAPI-negative cells was detected by flow cytometry (Fortessa, BD Biosciences) and analyzed using FlowJo analysis software (TreeStar). Heme/porphyrin analysis was analyzed by UPLC with an Acquity UPLC BEH C18, 1.7 μM, 2.1 × 100 mm column performed at the Iron and Heme Core facility of the University of Utah. Benzidine staining was performed by collecting 1×10^6^ K562 cells, washing with PBS, and staining with 0.2% benzidine dihydrochloride in 0.5% acetic acid for 15 minutes at room temperature. Benzidine positive cells were quantified by hemocytometer using bright-field microscopy. Experiments were performed in triplicate with 200 cells or more counted from randomly selected fields for each condition. Statistical significance was determined by two-sided unpaired *t*-tests using GraphPad Prism 7.0.

### Streptavidin affinity capture of biotin-modified proteins

To induce GFI1B-BirA*:HA in K562 cells, 6×10^7^ cells were treated with 1 μg/ml doxycycline for 48 hours. 30 μM biotin was added 16 hours before harvesting. Cells were collected, washed 2X with ice cold PBS, lysed with ice cold lysis buffer (50mM Tris pH7.5, 500 mM NaCl, 0.5 mM EDTA, 1% Triton X-100), sonicated and cleared by centrifugation. Biotinylated complexes were captured by SAv-Sepharose High Performance Beads and rocked at 4°C overnight. Beads were washed 5X with lysis buffer supplemented with SDS, boiled in DTT-containing sample buffer and fractionated by SDS-PAGE (10% T/2.67% C, 4% Stack). The gel was stained with Colloidal Coomassie Blue (Invitrogen). Lanes were divided into two equal sections for downstream LC-MS-MS analysis.

### Protein identification by mass spectrometry

Mass spectrometry was performed by the Taplin Mass Spectrometry Facility, Cell Biology Department, Harvard Medical School. Briefly, samples were reduced using 1 mM DTT in 50 mM ammonium bicarbonate for 30 minutes at 60°C, cooled and 5 mM iodoacetamide in 50 mM ammonium bicarbonate added for 15 minutes in the dark at ambient. 5 mM DTT was added to quench iodoacetamide derivatization. 5 ng/μl of sequencing-grad trypsin (Promega) was added and proteins digested overnight at 37°C. Samples were then desalted and reconstituted in 5-10 μl of HPLC solvent A (2.5% acetonitrile, 0.1% formic acid). A nano-scale reverse-phase HPLC capillary column was created by packing 2.6 μm C18 spherical silica beads into a fused silica capillary (100 μm inner diameter × ~30 cm length) with a flame-drawn tip. After column equilibration, each sample was loaded via a Famos auto sampler (LC Packings) onto the column. A linear gradient was formed to elute peptides with increasing concentrations of solvent B (97.5% acetonitrile, 0.1% formic acid). Eluted peptides were subjected to electrospray ionization coupled to an LTQ Orbitrap Velos Pro ion-trap mass spectrometer (Thermo-Fisher Scientific) to produce tandem mass spectra whose b-and y-ion series patterns were compared to theoretical digests of the proteome in Sequest (Thermo-Fisher Scientific) to establish protein identity. Data were filtered to a 1-2% peptide false discovery rate.

### Statistical analysis of mass spectrometry data

The read intensities were averaged across the biological triplicates for each sample. To establish the GFI1B proximitome (assessment of GFI1B-BirA* vs. BirA* only), sum read intensities among detected proteins were ranked from 0 to 100 percent. A total of 2751 proteins were detected in the BirA* only sample and were utilized to established a background intensity ranking. The intensity fold change for each ranked protein was determined by dividing the GFI1B-BirA*:HA average read intensity by the BirA*:HA only average read intensity. This calculated fold change of the intensity was compared to the average intensity rank and was observed to follow a strong diminution of variance with increasing values as expected. To identify proteins with aberrantly increased detection, z-scores were calculated for each of the 2751 proteins as the deviance in rank from the localized (across the background intensity axis) mean rank compared to the localized standard deviation of the ranks after the manner of Eubank (96). The corresponding p-value from a *t*-distribution of a 20-protein smoothing selection was determined and utilized to rank prioritize proteins showcasing the most consistent and increased abundance compared to expectation. Proteins with a p-value of <0.05 were considered hits. This approach was deemed appropriate as background intensity of pertinent proteins varied and detecting relative abundance compared to background was paramount. A volcano plot was created from the –log_10_ of the p-value and the log2 of the intensity fold change. Read intensities detected in GFI1B-BirA*:HA but not in the BirA*:HA only sample (500 proteins) were analyzed separately considering only the top 50-100 percentile of ranked GFI1B-BirA*:HA proximity partners as hits (a total of 35 proteins). Likewise, assessment of the GFI1B-BirA*:HA vs. GFI1B-ΔSNAG-BirA*:HA, GFI1B-BirA*:HA vs. GFI1B-P2A-BirA*:HA, and SNAG-BirA*:HA vs. BirA*:HA only was done by establishing a background percentile ranking of GFI1B-ΔSNAG-BirA*:HA (2897 proteins), GFI1B-P2A-BirA*:HA (2938 proteins), and BirA*:HA (2751 proteins). Proteins were then plotted according to background rank score and fold change. Statistical analysis was performed as described above for GFI1B-BirA*:HA vs. BirA*:HA using nonparametric regression and spline smoothing (97). SAS version 9.4 and Microsoft Excel were used for statistical calculations and data comparisons.

## ACKNOWLEDGEMENTS

The authors thank Katharine Ullman, Jason Gertz, Trudy Oliver, Charles Murtaugh, Rodney Stewart, and Mahesh Chandrasekharan for their helpful insights and suggestions during preparation of this manuscript. We acknowledge the following University of Utah Core facilities: DNA sequencing and cell line authentication was performed at the DNA Sequencing Core Facility. Oligonucleotides were synthesized by the DNA/Peptide Facility of the Health Sciences Center (HSC). Flow cytometry was performed by the Flow Cytometry Facility with support from the National Cancer Institute (5P30CA042014-24), and heme/porphyrin analysis was performed at the Heme & Iron Facility supported in part by a grant from the NIH National Institute of Diabetes and Digestive and Kidney Diseases, (U54DK110858). Mass spectrometry was performed at the Taplin Mass Spectrometry Facility, Cell Biology Department, Harvard Medical School with a special thanks to Ross Tomaino.

This work was supported by the National Institutes of Health (R01CA201235) (MEE) and T32 DK007115 (DM and MJC), and by grants from the American Cancer Society, Alex’s Lemonade Stand Foundation and the Hyundai Hope on Wheels Foundation. Supporters had no role in study design, data collection and interpretation, or the decision to submit the work for publication. The authors declare no conflicts of interest associated with the manuscript.

## SUPPLEMENTARY MATERIALS

**Supplementary Data S1** File: McClellan et al Mass Spec Data.xlsx (1.6 MB)

**Table S1.**
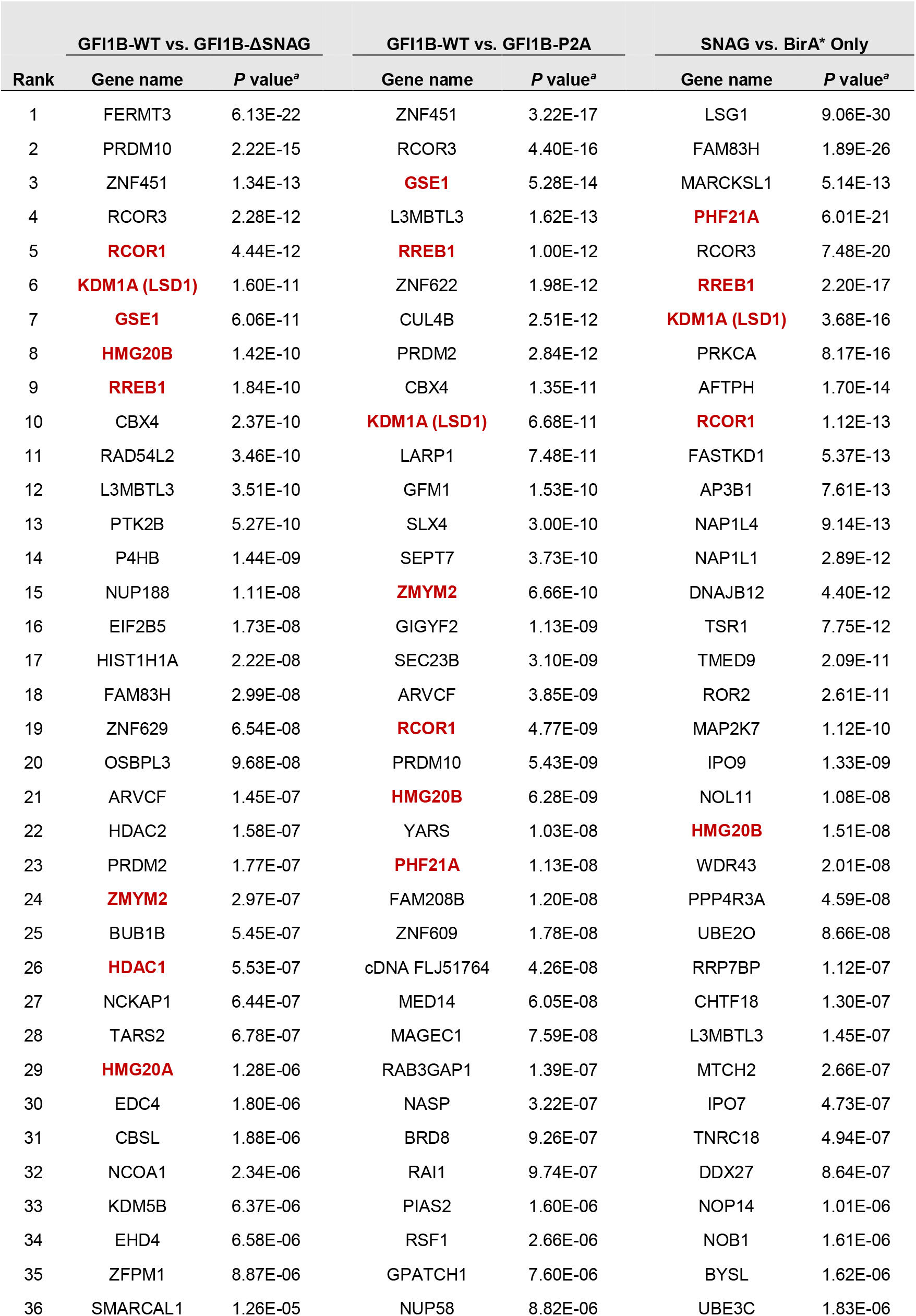
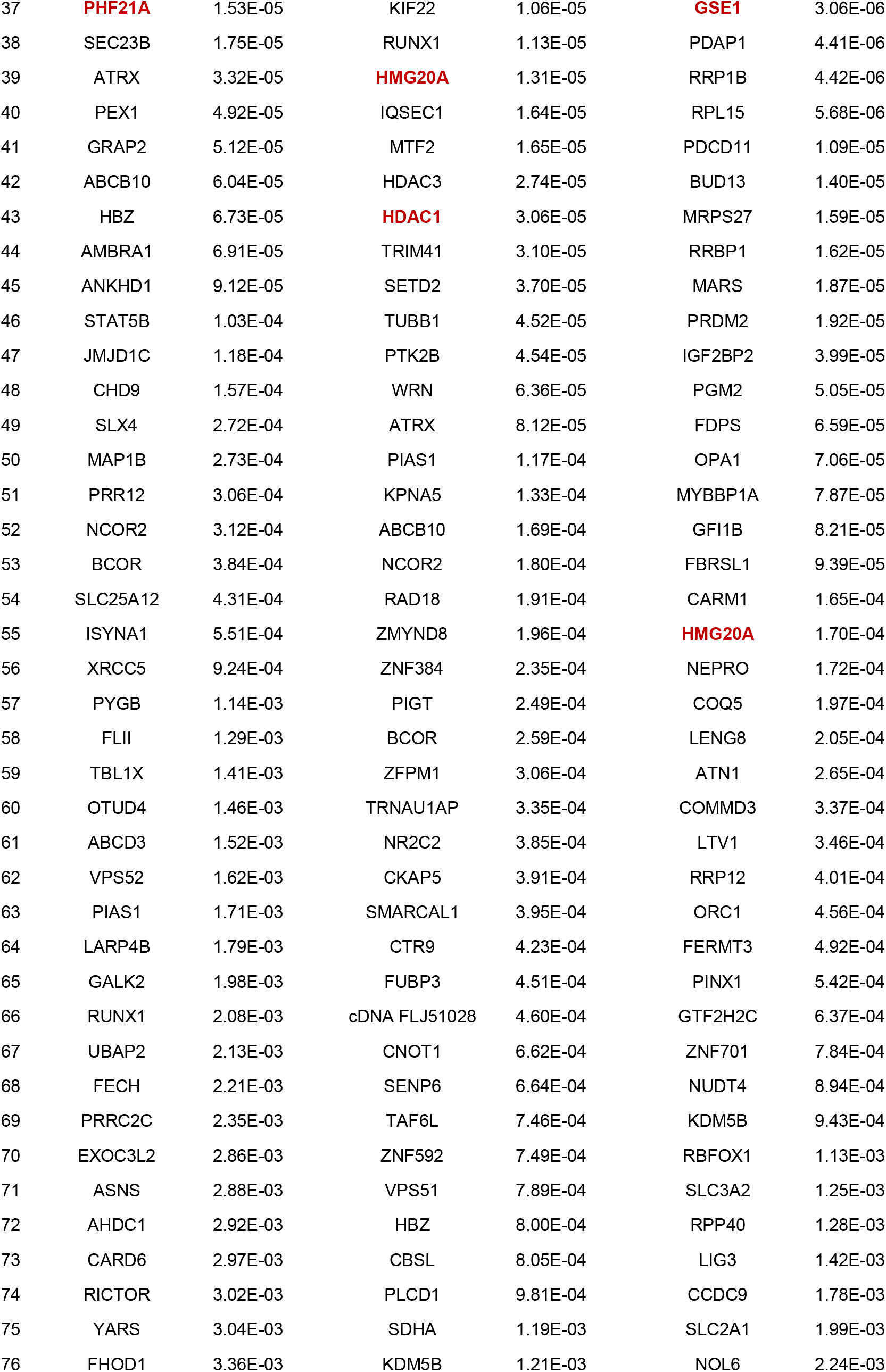
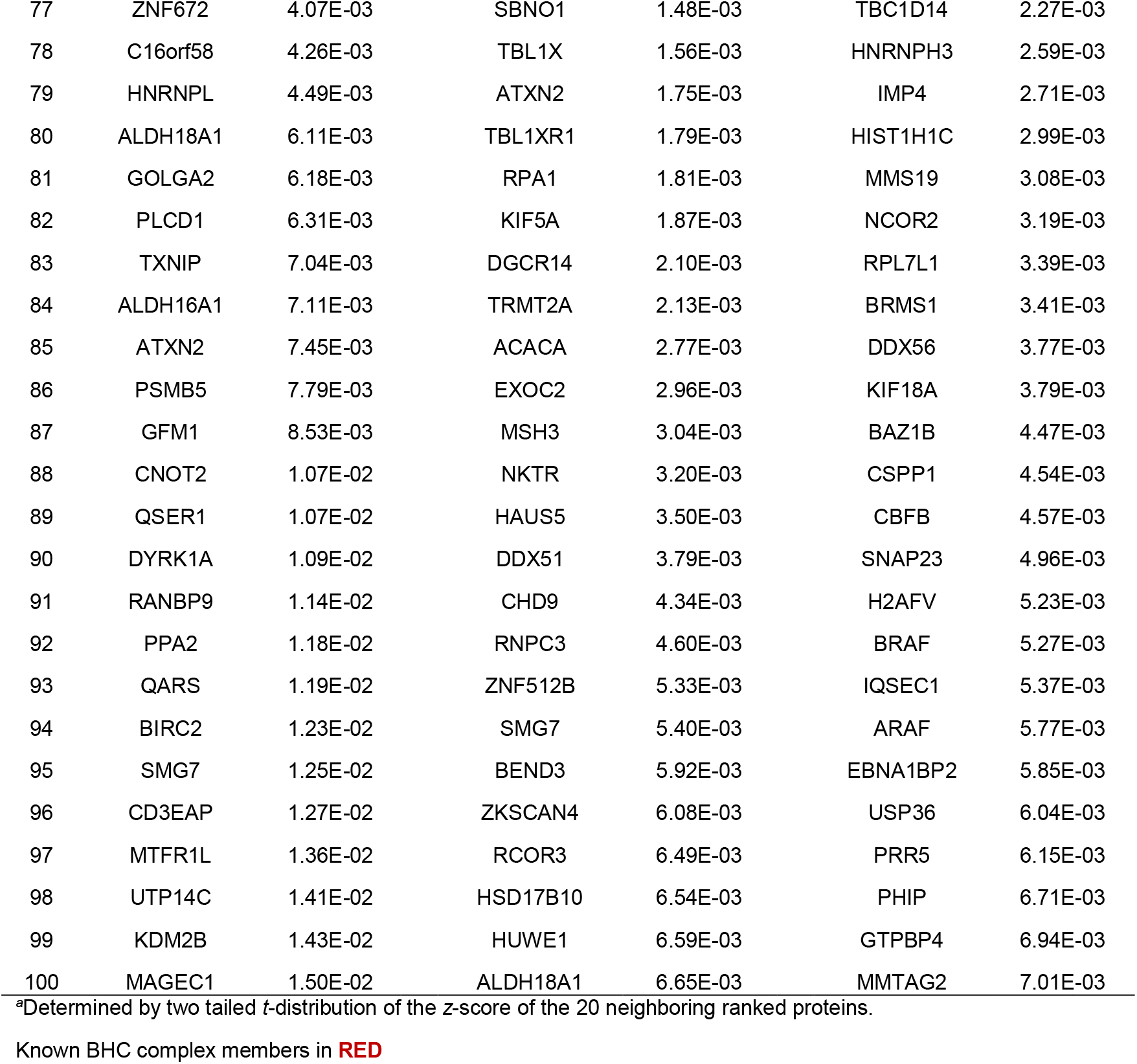
Top 100 LSD1-dependent proximity partners for each comparison

**Table S2.**
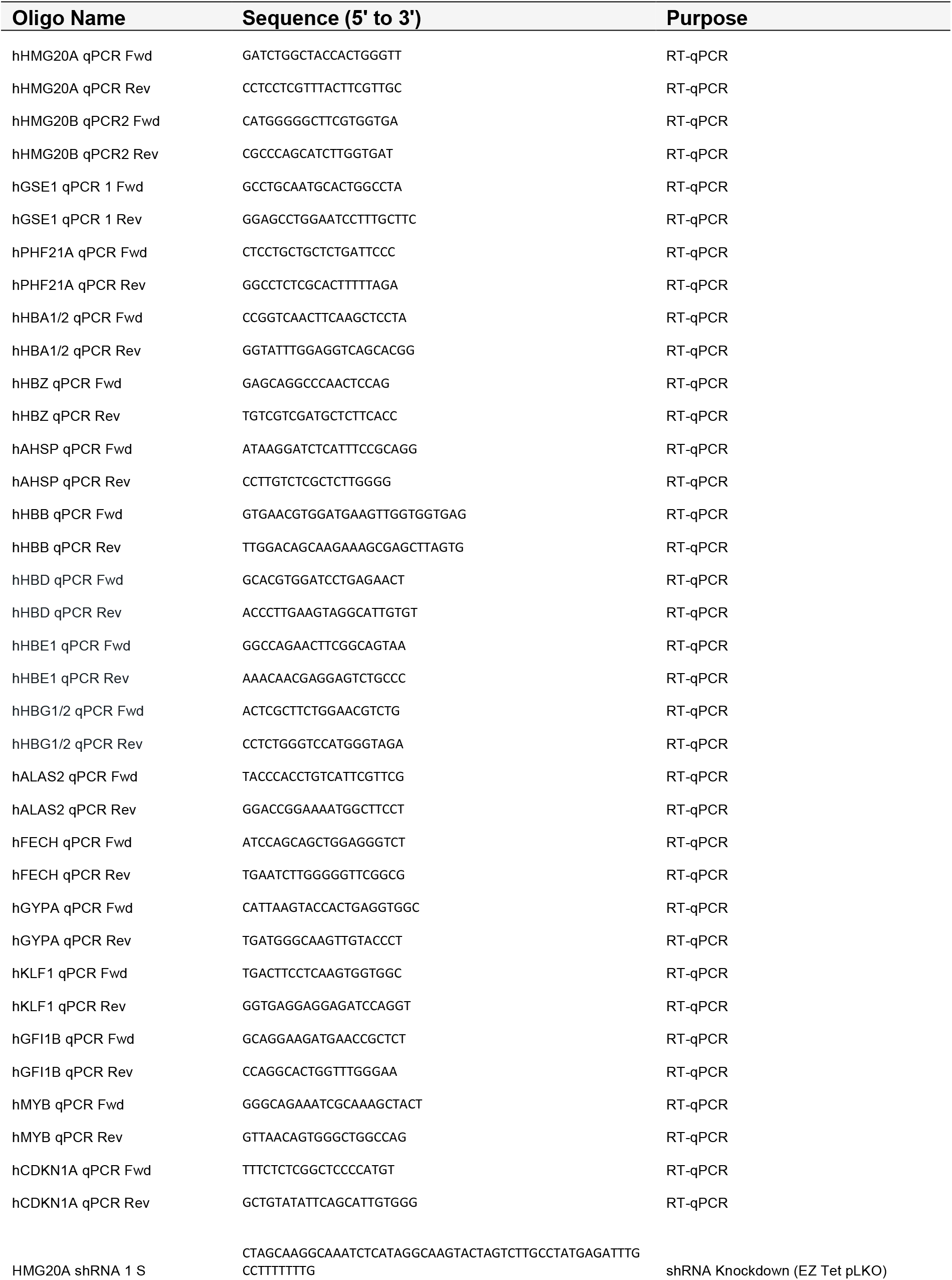
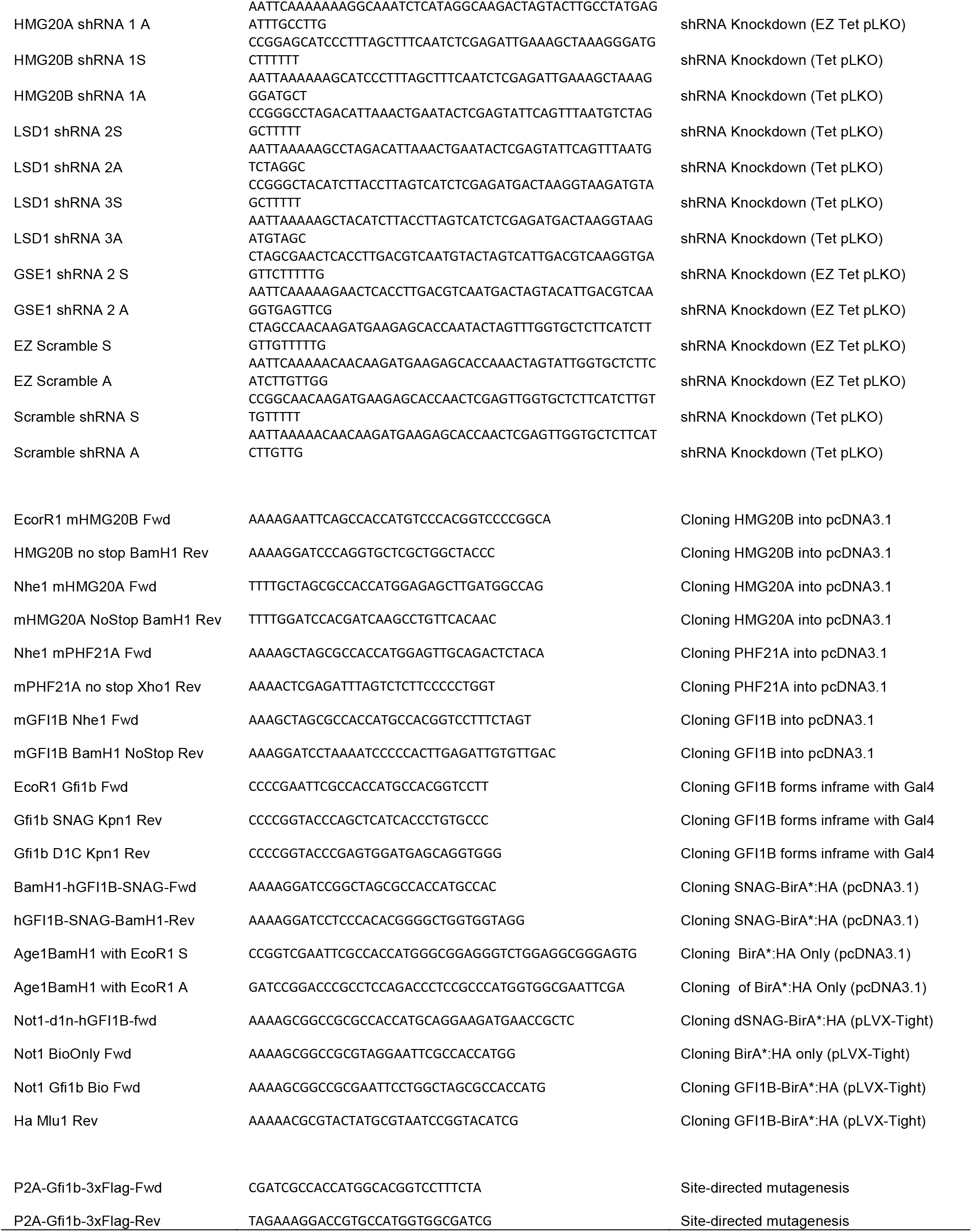
Oligonucleotides used in this study

## REFERENCES

1. Feinberg AP, Koldobskiy MA, Göndör A. 2016. Epigenetic modulators, modifiers and mediators in cancer aetiology and progression. Nat Rev Genet 17:284–299.

2. Avgustinova A, Benitah SA. 2016. Epigenetic control of adult stem cell function. Nat Rev Mol Cell Biol 17:643–658.

3. Ntziachristos P, Abdel-Wahab O, Aifantis I. 2016. Emerging concepts of epigenetic dysregulation in hematological malignancies. Nature Immunology 17:1016–1024.

4. Egger G, Liang G, Aparicio A, Jones PA. 2004. Epigenetics in human disease and prospects for epigenetic therapy. Nature 429:457–463.

5. van der Meer LT, Jansen JH, van der Reijden BA. 2010. Gfi1 and Gfi1b: key regulators of hematopoiesis. Leukemia 24:1834–1843.

6. Lancrin C, Mazan M, Stefanska M, Patel R, Lichtinger M, Costa G, Vargel O, Wilson NK, Möröy T, Bonifer C, Göttgens B, Kouskoff V, Lacaud G. 2012. GFI1 and GFI1B control the loss of endothelial identity of hemogenic endothelium during hematopoietic commitment. Blood 120:314–322.

7. Zeng H, Yücel R, Kosan C, Klein-Hitpass L, Möröy T. 2004. Transcription factor Gfi1 regulates self-renewal and engraftment of hematopoietic stem cells. EMBO J 23:4116–4125.

8. Hock H, Hamblen MJ, Rooke HM, Schindler JW, Saleque S, Fujiwara Y, Orkin SH. 2004. Gfi1 restricts proliferation and preserves functional integrity of haematopoietic stem cells. Nature 431:1002–1007.

9. Yücel R, Karsunky H, Klein-Hitpass L, Möröy T. 2003. The transcriptional repressor Gfi1 affects development of early, uncommitted c-Kit+ T cell progenitors and CD4/CD8 lineage decision in the thymus. J Exp Med 197:831–844.

10. Fiolka K, Hertzano R, Vassen L, Zeng H, Hermesh O, Avraham KB, Dührsen U, Möröy T. 2006. Gfi1 and Gfi1b act equivalently in haematopoiesis, but have distinct, non-overlapping functions in inner ear development. EMBO reports 7:326–333.

11. Karsunky H, Zeng H, Schmidt T, Zevnik B, Kluge R, Schmid KW, Dührsen U, Möröy T. 2002. Inflammatory reactions and severe neutropenia in mice lacking the transcriptional repressor Gfi1. Nat Genet 30:295–300.

12. Hock H, Hamblen MJ, Rooke HM, Traver D, Bronson RT, Cameron S, Orkin SH. 2003. Intrinsic requirement for zinc finger transcription factor Gfi-1 in neutrophil differentiation. Immunity 18:109–120.

13. Zörnig M, Schmidt T, Karsunky H, Grzeschiczek A, Moroy T. 1996. Zinc finger protein GFI-1 cooperates with myc and pim-1 in T-cell lymphomagenesis by reducing the requirements for IL-2. Oncogene 12:1789–1801.

14. Khandanpour C, Thiede C, Valk PJM, Sharif-Askari E, Nückel H, Lohmann D, Horsthemke B, Siffert W, Neubauer A, Grzeschik K-H, Bloomfield CD, Marcucci G, Maharry K, Slovak ML, Van der Reijden BA, Jansen JH, Schackert HK, Afshar K, Schnittger S, Peeters JK, Kroschinsky F, Ehninger G, Lowenberg B, Dührsen U, Möröy T. 2010. A variant allele of Growth Factor Independence 1 (GFI1) is associated with acute myeloid leukemia. Blood 115:2462–2472.

15. Khandanpour C. 2017. Growth factor independence 1 (Gfi1) regulates the AML supporting function of mesenchymal stromal cells. Experimental Hematology 53:S90.

16. Möröy T. 2014. The zinc finger protein Gfi1 maintains development and progression of lymphoid leukemia by blocking the activation of the tumor suppressor p53. Experimental Hematology 42:S7.

17. Volpe G, Walton DS, Grainger DE, Ward C, Cauchy P, Blakemore D, Coleman DJL, Cockerill PN, Garcia P, Frampton J. 2017. Prognostic significance of high GFI1 expression in AML of normal karyotype and its association with a FLT3-ITD signature. Sci Rep 7:11148.

18. Xia J, Bolyard AA, Rodger E, Stein S, Aprikyan AA, Dale DC, Link DC. 2009. Prevalence of mutations in ELANE, GFI1, HAX1, SBDS, WAS and G6PC3 in patients with severe congenital neutropenia. Br J Haematol 147:535–542.

19. Person RE, Li F-Q, Duan Z, Benson KF, Wechsler J, Papadaki HA, Eliopoulos G, Kaufman C, Bertolone SJ, Nakamoto B, Papayannopoulou T, Grimes HL, Horwitz M. 2003. Mutations in proto-oncogene GFI1 cause human neutropenia and target ELA2. Nat Genet 34:308–312.

20. Randrianarison-Huetz V, Laurent B, Bardet V, Blobe GC, Huetz F, Dumenil D. 2010. Gfi-1B controls human erythroid and megakaryocytic differentiation by regulating TGF-signaling at the bipotent erythro-megakaryocytic progenitor stage. Blood 115:2784–2795.

21. Vassen L, Beauchemin H, Lemsaddek W, Krongold J, Trudel M, Möröy T. 2014. Growth factor independence 1b (gfi1b) is important for the maturation of erythroid cells and the regulation of embryonic globin expression. PLoS ONE 9:e96636.

22. Singh D, Upadhyay G, Sengupta A, Biplob MA, Chakyayil S, George T, Saleque S. 2016. Cooperative Stimulation of Megakaryocytic Differentiation by Gfi1b Gene Targets Kindlin3 and Talin1. PLoS ONE 11:e0164506.

23. Beauchemin H, Shooshtarizadeh P, Vadnais C, Vassen L, Pastore YD, Möröy T. 2017. Gfi1b controls integrin signaling-dependent cytoskeleton dynamics and organization in megakaryocytes. Haematologica 102:484–497.

24. Saleque S, Cameron S, Orkin SH. 2002. The zinc-finger proto-oncogene Gfi-1b is essential for development of the erythroid and megakaryocytic lineages. Genes Dev 16:301–306.

25. Stevenson WS, Morel-Kopp M-C, Chen Q, Liang HP, Bromhead CJ, Wright S, Turakulov R, Ng AP, Roberts AW, Bahlo M, Ward CM. 2013. GFI1B mutation causes a bleeding disorder with abnormal platelet function. J Thromb Haemost 11:2039–2047.

26. Monteferrario D, Bolar NA, Marneth AE, Hebeda KM, Bergevoet SM, Veenstra H, Laros-van Gorkom BAP, MacKenzie MA, Khandanpour C, Botezatu L, Fransen E, Van Camp G, Duijnhouwer AL, Salemink S, Willemsen B, Huls G, Preijers F, Van Heerde W, Jansen JH, Kempers MJE, Loeys BL, Van Laer L, Van der Reijden BA. 2014. A dominant-negative GFI1B mutation in the gray platelet syndrome. N Engl J Med 370:245–253.

27. Uchiyama Y, Ogawa Y, Kunishima S, Shiina M, Nakashima M, Yanagisawa K, Yokohama A, Imagawa E, Miyatake S, Mizuguchi T, Takata A, Miyake N, Ogata K, Handa H, Matsumoto N. 2018. A novel GFI1B mutation at the first zinc finger domain causes congenital macrothrombocytopenia. Br J Haematol 181:843–847.

28. Kitamura K, Okuno Y, Yoshida K, Sanada M, Shiraishi Y, Muramatsu H, Kobayashi R, Furukawa K, Miyano S, Kojima S, Ogawa S, Kunishima S. 2016. Functional characterization of a novel GFI1B mutation causing congenital macrothrombocytopenia. J Thromb Haemost.

29. Vassen L, Khandanpour C, Ebeling P, Van der Reijden BA, Jansen JH, Mahlmann S, Dührsen U, Möröy T. 2009. Growth factor independent 1b (Gfi1b) and a new splice variant of Gfi1b are highly expressed in patients with acute and chronic leukemia. Int J Hematol 89:422–430.

30. Elmaagacli AH, Koldehoff M, Zakrzewski JL, Steckel NK, Ottinger H, Beelen DW. 2007. Growth factor-independent 1B gene (GFI1B) is overexpressed in erythropoietic and megakaryocytic malignancies and increases their proliferation rate. Br J Haematol 136:212–219.

31. Hernández A, Villegas A, Anguita E. 2010. Human promoter mutations unveil Oct-1 and GATA-1 opposite action on Gfi1b regulation. Annals of Hematology 89:759–765.

32. Tong B, Grimes HL, Yang TY, Bear SE, Qin Z, Du K, El-Deiry WS, Tsichlis PN. 1998. The Gfi-1B proto-oncoprotein represses p21WAF1 and inhibits myeloid cell differentiation. Mol Cell Biol 18:2462–2473.

33. Thivakaran A, Botezatu L, Hönes JM, Schütte J, Vassen L, Al-Matary YS, Patnana P, Zeller A, Heuser M, Thol F, Gabdoulline R, Olberding N, Frank D, Suslo M, Köster R, Lennartz K, Görgens A, Giebel B, Opalka B, Dührsen U, Khandanpour C. 2018. Gfi1b: a key player in the genesis and maintenance of acute myeloid leukemia and myelodysplastic syndrome. Haematologica 103:614–625.

34. Möröy T, Vassen L, Wilkes B, Khandanpour C. 2015. From cytopenia to leukemia: the role of Gfi1 and Gfi1b in blood formation. Blood 126:2561–2569.

35. Zweidler-McKay PA, Grimes HL, Flubacher MM, Tsichlis PN. 1996. Gfi-1 encodes a nuclear zinc finger protein that binds DNA and functions as a transcriptional repressor. Mol Cell Biol 16:4024–4034.

36. Lee S, Doddapaneni K, Hogue A, McGhee L, Meyers S, Wu Z. 2010. Solution structure of Gfi1 zinc domain bound to consensus DNA. J Mol Biol 397:1055–1066.

37. Anguita E, Villegas A, Iborra F, Hernández A. 2010. GFI1B controls its own expression binding to multiple sites. Haematologica 95:36–46.

38. Nakazawa Y, Suzuki M, Manabe N, Yamada T, Kihara-Negishi F, Sakurai T, Tenen DG, Iwama A, Mochizuki M, Oikawa T. 2007. Cooperative interaction between ETS1 and GFI1 transcription factors in the repression of Bax gene expression. Oncogene 26:3541–3550.

39. Huang D-Y, Kuo Y-Y, Chang Z-F. 2005. GATA-1 mediates auto-regulation of Gfi-1B transcription in K562 cells. Nucleic Acids Research 33:5331–5342.

40. Marteijn JAF, van der Meer LT, van Emst L, van Reijmersdal S, Wissink W, de Witte T, Jansen JH, Van der Reijden BA. 2007. Gfi1 ubiquitination and proteasomal degradation is inhibited by the ubiquitin ligase Triad1. Blood 110:3128–3135.

41. Dahl R, Iyer SR, Owens KS, Cuylear DD, Simon MC. 2007. The transcriptional repressor GFI1 antagonizes PU.1 activity through protein-protein interaction. J Biol Chem 282:6473–6483.

42. Andrade D, Velinder M, Singer J, Maese L, Bareyan D, Nguyen H, Chandrasekharan MB, Lucente H, McClellan D, Jones D, Sharma S, Liu F, Engel ME. 2016. SUMOylation Regulates Growth Factor Independence 1 in Transcriptional Control and Hematopoiesis. Mol Cell Biol 36:1438–1450.

43. Saleque S, Kim J, Rooke HM, Orkin SH. 2007. Epigenetic regulation of hematopoietic differentiation by Gfi-1 and Gfi-1b is mediated by the cofactors CoREST and LSD1. Mol Cell 27:562–572.

44. Lin Y, Wu Y, Li J, Dong C, Ye X, Chi Y-I, Evers BM, Zhou BP. 2010. The SNAG domain of Snail1 functions as a molecular hook for recruiting lysine-specific demethylase 1. EMBO J 29:1803–1816.

45. Laurent B, Randrianarison-Huetz V, Frisan E, Andrieu-Soler C, Soler E, Fontenay M, Dusanter-Fourt I, Duménil D. 2012. A short Gfi-1B isoform controls erythroid differentiation by recruiting the LSD1-CoREST complex through the dimethylation of its SNAG domain. J Cell Sci 125:993–1002.

46. Velinder M, Singer J, Bareyan D, Meznarich J, Tracy CM, Fulcher JM, McClellan D, Lucente H, Franklin S, Sharma S, Engel ME. 2017. GFI1 functions in transcriptional control and cell fate determination require SNAG domain methylation to recruit LSD1. Biochem J 474:2951–2951.

47. Upadhyay G, Chowdhury AH, Vaidyanathan B, Kim D, Saleque S. 2014. Antagonistic actions of Rcor proteins regulate LSD1 activity and cellular differentiation. Proc Natl Acad Sci USA 111:8071–8076.

48. Shi Y, Lan F, Matson C, Mulligan P, Whetstine JR, Cole PA, Casero RA, Shi Y. 2004. Histone demethylation mediated by the nuclear amine oxidase homolog LSD1. Cell 119:941–953.

49. Metzger E, Wissmann M, Yin N, Müller JM, Schneider R, Peters AHFM, Günther T, Buettner R, Schüle R. 2005. LSD1 demethylates repressive histone marks to promote androgen-receptor-dependent transcription. Nature 437:436–439.

50. Huang J, Sengupta R, Espejo AB, Lee MG, Dorsey JA, Richter M, Opravil S, Shiekhattar R, Bedford MT, Jenuwein T, Berger SL. 2007. p53 is regulated by the lysine demethylase LSD1. Nature 449:105–108.

51. Yang J, Huang J, Dasgupta M, Sears N, Miyagi M, Wang B, Chance MR, Chen X, Du Y, Wang Y, An L, Wang Q, Lu T, Zhang X, Wang Z, Stark GR. 2010. Reversible methylation of promoter-bound STAT3 by histone-modifying enzymes. Proc Natl Acad Sci USA 107:21499–21504.

52. Zhang X, Tanaka K, Yan J, Li J, Peng D, Jiang Y, Yang Z, Barton MC, Wen H, Shi X. 2013. Regulation of estrogen receptor α by histone methyltransferase SMYD2-mediated protein methylation. Proc Natl Acad Sci USA 110:17284–17289.

53. Cho H-S, Suzuki T, Dohmae N, Hayami S, Unoki M, Yoshimatsu M, Toyokawa G, Takawa M, Chen T, Kurash JK, Field HI, Ponder BAJ, Nakamura Y, Hamamoto R. 2011. Demethylation of RB regulator MYPT1 by histone demethylase LSD1 promotes cell cycle progression in cancer cells. Cancer Res 71:655–660.

54. Lee J-Y, Park J-H, Choi H-J, Won H-Y, Joo H-S, Shin D-H, Park MK, Han B, Kim KP, Lee TJ, Croce CM, Kong G. 2017. LSD1 demethylates HIF1α to inhibit hydroxylation and ubiquitin-mediated degradation in tumor angiogenesis. Oncogene 36:5512–5521.

55. Wang J, Hevi S, Kurash JK, Lei H, Gay F, Bajko J, Su H, Sun W, Chang H, Xu G, Gaudet F, Li E, Chen T. 2009. The lysine demethylase LSD1 (KDM1) is required for maintenance of global DNA methylation. Nat Genet 41:125–129.

56. Kontaki H, Talianidis I. 2010. Lysine methylation regulates E2F1-induced cell death. Mol Cell 39:152–160.

57. Zhang C, Hoang N, Leng F, Saxena L, Lee L, Alejo S, Qi D, Khal A, Sun H, Lu F, Zhang H. 2018. LSD1 demethylase and the methyl-binding protein PHF20L1 prevent SET7 methyltransferase-dependent proteolysis of the stem-cell protein SOX2. J Biol Chem 293:3663–3674.

58. Wang Y, Zhang H, Chen Y, Sun Y, Yang F, Yu W, Liang J, Sun L, Yang X, Shi L, Li R, Li Y, Zhang Y, Li Q, Yi X, Shang Y. 2009. LSD1 is a subunit of the NuRD complex and targets the metastasis programs in breast cancer. Cell 138:660–672.

59. Yang Y, Huang W, Qiu R, Liu R, Zeng Y, Gao J, Zheng Y, Hou Y, Wang S, Yu W, Leng S, Feng D, Wang Y. 2018. LSD1 coordinates with the SIN3A/HDAC complex and maintains sensitivity to chemotherapy in breast cancer. J Mol Cell Biol 387:49.

60. Ray SK, Li HJ, Metzger E, Schüle R, Leiter AB. 2014. CtBP and associated LSD1 are required for transcriptional activation by NeuroD1 in gastrointestinal endocrine cells. Mol Cell Biol 34:2308–2317.

61. Tsai M-C, Manor O, Wan Y, Mosammaparast N, Wang JK, Lan F, Shi Y, Segal E, Chang HY. 2010. Long noncoding RNA as modular scaffold of histone modification complexes. Science 329:689–693.

62. Shi Y-J, Matson C, Lan F, Iwase S, Baba T, Shi Y. 2005. Regulation of LSD1 histone demethylase activity by its associated factors. Mol Cell 19:857–864.

63. Roux KJ, Kim DI, Raida M, Burke B. 2012. A promiscuous biotin ligase fusion protein identifies proximal and interacting proteins in mammalian cells. J Cell Biol 196:801–810.

64. Kim DI, Jensen SC, Noble KA, Kc B, Roux KH, Motamedchaboki K, Roux KJ. 2016. An improved smaller biotin ligase for BioID proximity labeling. Mol Biol Cell 27:1188–1196.

65. Hakimi M-A, Bochar DA, Chenoweth J, Lane WS, Mandel G, Shiekhattar R. 2002. A core-BRAF35 complex containing histone deacetylase mediates repression of neuronal-specific genes. Proc Natl Acad Sci USA 99:7420–7425.

66. Gocke CB, Yu H. 2008. ZNF198 stabilizes the LSD1-CoREST-HDAC1 complex on chromatin through its MYM-type zinc fingers. PLoS ONE 3:e3255.

67. Rivero S, Ceballos-Chávez M, Bhattacharya SS, Reyes JC. 2015. HMG20A is required for SNAI1-mediated epithelial to mesenchymal transition. Oncogene 34:5264–5276.

68. Garçon L, Lacout C, Svinartchouk F, Le Couédic J-P, Villeval J-L, Vainchenker W, Duménil D. 2005. Gfi-1B plays a critical role in terminal differentiation of normal and transformed erythroid progenitor cells. Blood 105:1448–1455.

69. Lam LT, Ronchini C, Norton J, Capobianco AJ, Bresnick EH. 2000. Suppression of erythroid but not megakaryocytic differentiation of human K562 erythroleukemic cells by notch-1. J Biol Chem 275:19676–19684.

70. Meng Y-S, Khoury H, Dick JE, Minden MD. 2005. Oncogenic potential of the transcription factor LYL1 in acute myeloblastic leukemia. Leukemia 19:1941–1947.

71. Maiques-Diaz A, Somervaille TC. 2016. LSD1: biologic roles and therapeutic targeting. Epigenomics 8:1103–1116.

72. Kim DI, Roux KJ. 2016. Filling the Void: Proximity-Based Labeling of Proteins in Living Cells. Trends Cell Biol 26:804–817.

73. Leon-Del-Rio A, Gravel RA. 1994. Sequence requirements for the biotinylation of carboxyl-terminal fragments of human propionyl-CoA carboxylase alpha subunit expressed in Escherichia coli. J Biol Chem 269:22964–22968.

74. Vassen L. 2005. Direct transcriptional repression of the genes encoding the zinc-finger proteins Gfi1b and Gfi1 by Gfi1b. Nucleic Acids Research 33:987–998.

75. Rodriguez P, Bonte E, Krijgsveld J, Kolodziej KE, Guyot B, Heck AJR, Vyas P, de Boer E, Grosveld F, Strouboulis J. 2005. GATA-1 forms distinct activating and repressive complexes in erythroid cells. EMBO J 24:2354–2366.

76. Kuo Y-Y, Chang Z-F. 2007. GATA-1 and Gfi-1B interplay to regulate Bcl-xL transcription. Mol Cell Biol 27:4261–4272.

77. Jegalian AG, Wu H. 2002. Regulation of Socs gene expression by the proto-oncoprotein GFI-1B: two routes for STAT5 target gene induction by erythropoietin. J Biol Chem 277:2345–2352.

78. Chowdhury AH, Ramroop JR, Upadhyay G, Sengupta A, Andrzejczyk A, Saleque S. 2013. Differential transcriptional regulation of meis1 by Gfi1b and its co-factors LSD1 and CoREST. PLoS ONE 8:e53666.

79. Vassen L, Fiolka K, Möröy T. 2006. Gfi1b alters histone methylation at target gene promoters and sites of gamma-satellite containing heterochromatin. EMBO J 25:2409–2419.

80. Ceballos-Chávez M, Rivero S, García-Gutiérrez P, Rodríguez-Paredes M, García-Domínguez M, Bhattacharya S, Reyes JC. 2012. Control of neuronal differentiation by sumoylation of BRAF35, a subunit of the LSD1-CoREST histone demethylase complex. Proc Natl Acad Sci USA 109:8085–8090.

81. Esteghamat F, van Dijk TB, Braun H, Dekker S, van der Linden R, Hou J, Fanis P, Demmers J, van IJcken W, Ozgür Z, Horos R, Pourfarzad F, Lindern von M, Philipsen S. 2011. The DNA binding factor Hmg20b is a repressor of erythroid differentiation. Haematologica 96:1252–1260.

82. Hock R, Furusawa T, Ueda T, Bustin M. 2007. HMG chromosomal proteins in development and disease. Trends Cell Biol 17:72–79.

83. Malarkey CS, Churchill MEA. 2012. The high mobility group box: the ultimate utility player of a cell. Trends Biochem Sci 37:553–562.

84. Wynder C, Hakimi M-A, Epstein JA, Shilatifard A, Shiekhattar R. 2005. Recruitment of MLL by HMG-domain protein iBRAF promotes neural differentiation. Nat Cell Biol 7:1113–1117.

85. Chai P, Tian J, Zhao D, Zhang H, Cui J, Ding K, Liu B. 2016. GSE1 negative regulation by miR-489-5p promotes breast cancer cell proliferation and invasion. Biochem Biophys Res Commun 471:123–128.

86. Ding K, Tan S, Huang X, Wang X, Li X, Fan R, Zhu Y, Lobie PE, Wang W, Wu Z. 2018. GSE1 predicts poor survival outcome in gastric cancer patients by SLC7A5 enhancement of tumor growth and metastasis. J Biol Chem 293:3949–3964.

87. Lambert SA, Jolma A, Campitelli LF, Das PK, Yin Y, Albu M, Chen X, Taipale J, Hughes TR, Weirauch MT. 2018. The Human Transcription Factors. Cell 172:650–665.

88. Clapier CR, Cairns BR. 2009. The biology of chromatin remodeling complexes. Annu Rev Biochem 78:273–304.

89. Grimes HL, Chan TO, Zweidler-McKay PA, Tong B, Tsichlis PN. 1996. The Gfi-1 proto-oncoprotein contains a novel transcriptional repressor domain, SNAG, and inhibits G1 arrest induced by interleukin-2 withdrawal. Mol Cell Biol 16:6263–6272.

90. Chen Y, Yang Y, Wang F, Wan K, Yamane K, Zhang Y, Lei M. 2006. Crystal structure of human histone lysine-specific demethylase 1 (LSD1). Proc Natl Acad Sci USA 103:13956–13961.

91. Yang M, Gocke CB, Luo X, Borek D, Tomchick DR, Machius M, Otwinowski Z, Yu H. 2006. Structural basis for CoREST-dependent demethylation of nucleosomes by the human LSD1 histone demethylase. Mol Cell 23:377–387.

92. Ouyang J, Shi Y, Valin A, Xuan Y, Gill G. 2009. Direct binding of CoREST1 to SUMO-2/3 contributes to gene-specific repression by the LSD1/CoREST1/HDAC complex. Mol Cell 34:145–154.

93. Nalawansha DA, Pflum MKH. 2017. LSD1 Substrate Binding and Gene Expression Are Affected by HDAC1-Mediated Deacetylation. ACS Chem Biol 12:254–264.

94. Holmes A, Roseaulin L, Schurra C, Waxin H, Lambert S, Zaratiegui M, Martienssen RA, Arcangioli B. 2012. Lsd1 and lsd2 control programmed replication fork pauses and imprinting in fission yeast. Cell Rep 2:1513–1520.

95. Sumoy L, Carim L, Escarceller M, Nadal M, Gratacòs M, Pujana MA, Estivill X, Peral B. 2000. HMG20A and HMG20B map to human chromosomes 15q24 and 19p13.3 and constitute a distinct class of HMG-box genes with ubiquitous expression. Cytogenet Cell Genet 88:62–67.

96. Eubank RL. 1999. Nonparametric Regression and Spline Smoothing, Second Edition. CRC Press.

97. Ke C, Wang Y. 2004. Smoothing Spline Nonlinear Nonparametric Regression Models. Journal of the American Statistical Association 99:1166–1175.

